# Carnosine synthase deficiency aggravates neuroinflammation in multiple sclerosis

**DOI:** 10.1101/2023.03.30.534899

**Authors:** Jan Spaas, Thibaux Van der Stede, Sarah de Jager, Annet van de Waterweg Berends, Assia Tiane, Hans Baelde, Shahid P. Baba, Matthias Eckhardt, Esther Wolfs, Tim Vanmierlo, Niels Hellings, Bert O. Eijnde, Wim Derave

## Abstract

Multiple sclerosis (MS) pathology features autoimmune-driven neuroinflammation, demyelination, and failed remyelination. Carnosine is a histidine-containing dipeptide (HCD) with pluripotent homeostatic properties that is able to improve outcomes in an animal MS model (EAE) when supplied exogenously. To uncover if endogenous carnosine is involved in, and protects against, MS-related neuroinflammation, demyelination or remyelination failure, we here studied the HCD-synthesizing enzyme carnosine synthase (CARNS1) in human MS lesions and two preclinical mouse MS models (EAE, cuprizone). We demonstrate that due to its presence in oligodendrocytes, CARNS1 expression is diminished in demyelinated MS lesions and mouse models mimicking demyelination/inflammation, but returns upon remyelination. *Carns1*-KO mice that are devoid of endogenous HCDs display exaggerated neuroinflammation and clinical symptoms during EAE, which could be partially rescued by exogenous carnosine treatment. Worsening of the disease appears to be driven by a central, not peripheral immune-modulatory, mechanism possibly linked to impaired clearance of the reactive carbonyl acrolein in *Carns1*-KO mice. In contrast, the presence of CARNS1 and endogenous HCDs does not protect against cuprizone-induced demyelination, and is not required for normal oligodendrocyte precursor cell differentiation and (re)myelin to occur. Exogenously administered carnosine is not effective in blunting demyelination or accelerating remyelination. In conclusion, we show that CARNS1 is diminished in demyelinated MS lesions, which may have detrimental effects on disease progression through weakening the endogenous protection against neuroinflammation.

## Introduction

Multiple sclerosis (MS) is an autoimmune disease that triggers multifocal inflammation and demyelination of the central nervous system (CNS) [1]. Neurodegeneration already starts to develop early in the disease course and becomes more pronounced in later stages, especially as remyelination fails [2, 3]. The last 30 years have seen dramatic progress in our understanding and treatment of the early, immune-mediated relapsing-remitting disease stage. Peripheral immune-modulatory therapies, however, fail to impact the later stages of the disease; suggesting distinct pathophysiological mechanisms [1, 4]. Indeed, immune cell infiltration into the CNS eventually wanes, but sustained CNS-compartmentalized inflammation and remyelination failure remain [5]. To accelerate the development of therapies that prevent MS progression or promote myelin repair, a better understanding of CNS-intrinsic pathophysiology is imperative.

Oxidative stress is one of the dominant pathological hallmarks that accompanies neuroinflammation and drives disease progression in MS [6]. It is linked to greater pro-inflammatory activity and myelin damage [7, 8], early and long-term axonal damage [9, 10], and impaired remyelination [11]. We recently reported that inflammation-mediated oxidative stress results in the formation of acrolein, a lipid-derived reactive carbonyl with pro-inflammatory and cytotoxic properties, in CNS lesions of MS patients and experimental autoimmune encephalomyelitis (EAE) mice [12]. In parallel, EAE mice displayed reduced levels of the dipeptide carnosine (β-alanyl-L-histidine), an endogenous carbonyl quencher with particularly high acrolein reactivity [13, 14]. A nutrition-induced increase of CNS carnosine levels amplified carnosine-acrolein adduct formation, while suppressing chronic (glial) inflammatory activity, acrolein-protein damage, and clinical disease severity in EAE [12].

Carnosine is the archetype of a larger family of histidine-containing dipeptides (HCDs), which also comprises homocarnosine (γ-aminobutyryl-L-histidine) and several less abundant methylated or acetylated carnosine analogs (anserine, balenine, N-acetylcarnosine) [15]. The enzyme carnosine synthase (CARNS1), an ATP-dependent ligase from the ‘ATP-grasp family’, accounts for the endogenous synthesis of (homo)carnosine [16]. RNA sequencing (e.g. [17, 18]) and immunostaining [19] studies indicate that mature oligodendrocytes are the major source of CARNS1 within the CNS. Other glia and neurons express transporters and enzymes to take up, release, and/or hydrolyse (homo)carnosine [17, 20–22]; implying that HCD metabolism is quite widely dispersed in the CNS. Yet, the physiological functions of HCDs in the CNS remain elusive, as is their role in the protection against neurological disease. In addition to our finding of a disturbed endogenous HCD homeostasis in EAE mice [12], recent transcriptome analyses suggest an altered, mostly reduced, *CARNS1* expression in demyelinating pathologies including MS [23–26]. In this manuscript, we hypothesized that the MS-induced loss of oligodendrocytes reduces CARNS1 levels and HCD synthesis, culminating in a weakened protection against neuroinflammation-related processes and an impaired remyelination potential.

## Materials and methods

### Immunohistochemistry on human brain

Post-mortem frozen brain tissue from MS patients and healthy controls was obtained from The Netherlands Brain Bank (NBB). Clinical details are depicted in **Supplementary Table S1**. Lesions were identified by proteolipid protein (PLP) and CD68 or HLA-DR staining to assess demyelination and the presence and distribution of microglia/macrophages, respectively [27]. To visualise CARNS1, sections (10 µm) were fixated with acetone (10 min), blocked with 10% donkey serum in PBS with 1% bovine serum albumin (BSA, 30 min), and exposed overnight at 4°C to primary CARNS1 antibodies (**Supplementary Table S2**). Hereafter, sections were washed and exposed to complementary secondary antibodies for 60 min (Thermo Fisher). In case of excessive background staining, 0.1-0.3% sudan black was applied (10 min). Fluorescence imaging was performed with a Leica DM4000 B LED (Leica Microsystems). For double-labelling, the following cell type markers were stained: OLIG2, CD68, glial fibrillary acidic protein (GFAP), and neurofilament heavy polypeptide (NF-H); see **Supplementary Table S2**.

### Gene expression in human brain

To assess *CARNS1* gene expression, total RNA was isolated from MS lesion tissue, the surrounding normal-appearing white matter (NAWM) and white matter tissue from matched healthy controls using the RNeasy mini kit (Qiagen). For this purpose, we selected chronic non-fibrotic demyelinated (PLP^−^) MS lesions from white matter (NeuN^−^) tissue, that did not contain HLA-DR^+^ or Oil Red O (ORO)^+^ immune cells on immunohistochemistry [28, 29]. Patient details are depicted in **Supplementary Table S3**. RNA concentration was measured with a NanoDrop spectrophotometer (Isogen Life Science). cDNA was synthesized by reverse-transcription using qScript cDNA Supermix kit (Quanta). Quantitative PCR (qPCR) was conducted on a Biosystems QuantStudio 3 system (Life Technologies) using 12.5 ng cDNA template, Fast SYBR Green Master Mix (Applied Biosystems), and the appropriate primer pairs (**Supplementary Table S4**). Data was analysed by the 2^−ΔΔCt^ method, normalised to the most stable reference genes (geNorm [30]), and expressed as fold changes.

### UHPLC-MS/MS on human plasma and cerebrospinal fluid

Blood and cerebrospinal fluid (CSF) samples were obtained via the University Biobank Limburg (UbiLim), with approval from the Medical Ethics Committee at Hasselt University. All participants signed informed consent. Venous blood samples from people with MS and age-matched healthy donors (2:1 ratio) were collected in EDTA-coated tubes, and plasma was stored at −80°C after centrifugation. Lumbar CSF from people with and without the diagnosis of MS was collected into PPS tubes, kept at 4°C, centrifuged to remove cells (500 × g, 10 min, 4°C), and supernatant was stored at −80°C. Clinical detail of all participants involved in blood and CSF sampling are shown in **Supplementary Table S5**. To determine HCD levels in human plasma and CSF by UHPLC-MS/MS (as described previously [19, 31]), deproteinized plasma or CSF (150 µL) was diluted in acetonitrile (215 µL), 1 µM carnosine-d4 as internal standard (10 µL), and ultrapure water (25 µL). Samples were thoroughly vortexed and centrifuged (15 min, 15000 × g, 4°C). A volume of 2.5 µL from the supernatant was injected in a Xevo TQ-S MS/MS® system. Standard calibration curves with all HCDs were prepared either in ultrapure water (for CSF), or in a pool of human deproteinized plasma that was collected after a 2-day lacto-ovo vegetarian diet to minimize circulating HCDs and thereby account for potential matrix effects during UHPLC-MS/MS.

### CN1 activity in human plasma

The CN1 activity was quantified in plasma from people with MS and age-matched controls (**Supplementary Table S6**). Blood was collected in heparin-coated tubes, centrifuged, and frozen immediately. Plasma samples were exposed to 10 mM carnosine (Flamma) at 37°C for 10 min, after which CN1 was stopped by adding 600 mM trichloroacetic acid (TCA). For controls, TCA was added before carnosine. Following centrifugation (15 min, 16000 × g), supernatants were incubated in a mixture of OPA (incomplete opthaldehyde with 0.2% 2-mercaptoethanol) and 4M sodium hydroxide for 40 min. Fluorimetric determination of liberated histidine was performed at 360 nm excitation wavelength, 465 nm emission wavelength (Infinite 200 PRO). Concentrations were calculated from a histidine standard curve. Results were expressed in µmol/mL/h.

### *Carns1*-KO mice

Whole-body *Carns1*-KO mice with a C57BL/6J background, recently developed by Wang-Eckhardt *et al.*, lack the CARNS1 protein and are devoid of endogenous HCDs (see [32], confirmed by our laboratory using highly sensitive UHPLC-MS/MS [19]. Mice were housed under standard room conditions (12h:12h light:dark cycle, 20-24°C, relative humidity 30-60%) and had *ad libitum* access to drinking water and food pellets. Following in-house breeding of *Carns1*-KO and WT littermates from heterozygous parents, female offspring was selected for EAE experiments whilst males were used for cuprizone experiments. Genotyping was performed in toe samples at ∼P10 using a previously published protocol [32]. All procedures were approved by the Ethical Committee on Animal Experiments at Hasselt University.

### EAE experiments

EAE was induced in 9-13 week old *Carns1*-KO and WT mice by active immunization using two subcutaneous injections (upper and lower back) of 100 µL myelin oligodendrocyte glycoprotein peptide fragment 35-55 (MOG_35-55_) emulsifed in complete Freund’s adjuvant (CFA). Intraperitoneal (i.p.) injections of pertussis toxin (110 ng in 100 µL PBS) were administered immediately hereafter and 24 h later (EK-2110, Hooke Laboratories). Body weight and clinical symptoms were monitored daily (scale 0-5 with 0.5 increments: 0, no symptoms; 1, limp tail; 2, hindlimb paresis; 3, hindlimb paralysis; 4, front limb paresis; 5, death due to EAE). Healthy control mice were included as well.

Part of the *Carns1*-KO mice received 30 g/L (3%) L-Carnosine (Flamma) in the drinking water starting 7 days prior to EAE induction; which attenuates EAE severity in WT mice [12]. EAE mice were sacrificed at the disease peak (17 days post-immunization) or in the chronic stage (day 46-56). Following overdose injection of Dolethal (200 mg/kg, i.p.), mice were perfused via a left ventricular puncture with 0.9% NaCl solution containing heparin (25 UI/mL). Spinal cord segments were isolated and either frozen directly and kept at −80°C (for qPCR, immunohistochemistry, western blot, UHPLC-MS/MS), or pooled with the brain and kept on ice in RPMI-1640 medium (Gibco) with 0.5% Pen-Strep (for flow cytometry). Spleens and lymph nodes were dissected at disease peak and processed for flow cytometry as well.

### Cuprizone experiments

Cuprizone (0.3% w/w, C9012, Sigma) was mixed with powdered chow and fed to 8-10 week old *Carns1*-KO and WT mice for 6 weeks to evoke acute demyelination. Hereafter, mice were sacrificed either immediately or following a 1-week recovery period with normal powdered chow to allow remyelination. Healthy control mice received powdered chow without cuprizone for 6 weeks. Body weight was monitored every second day. Brains were isolated from mice following Dolethal overdose and perfusion, as described above. The corpus callosum was dissected at approximately Bregma level 0 to −1 mm, frozen immediately and stored at −80°C (for UHPLC-MS/MS, western blot), or fixated in 2% glutaraldehyde and kept at 4°C (for electron microscopy). The dorsal part of the brain was prepared for immunohistochemistry by either gradually freezing on a dry ice-cooled metal plate, or by overnight fixation in 4% paraformaldehyde (PFA) followed by 10-20-30% sucrose series. Next, brains were stored at −80°C until analysis.

In a separate experiment, 8-week old WT male C57BL/6J mice were obtained from Envigo (The Netherlands) and subjected to a 0.3% cuprizone diet for 6 weeks, with or without 3% carnosine added to the drinking water. Healthy control mice received normal powdered chow for 6 weeks. In addition, to study the effect of exogenous carnosine on remyelination, mice received normal tap water or tap water containing 3% carnosine during the last week of the cuprizone feeding and during the 1-week recovery phase. An extra control group, receiving 3% carnosine during the last week of cuprizone feeding and sacrificed before the recovery week started, was added to exclude any effects during this ‘loading phase’. Body weight was monitored daily. Tissue dissection and processing were performed as described above, except that brain tissue for immunohistochemistry was fixated in 4% PFA (24 h), kept in 70% ethanol, and embedded in paraffin.

To measure *Carns1* mRNA at multiple time points (see **Fig 1i**), samples from a previous cuprizone study were used [33].

**Fig. 1.**
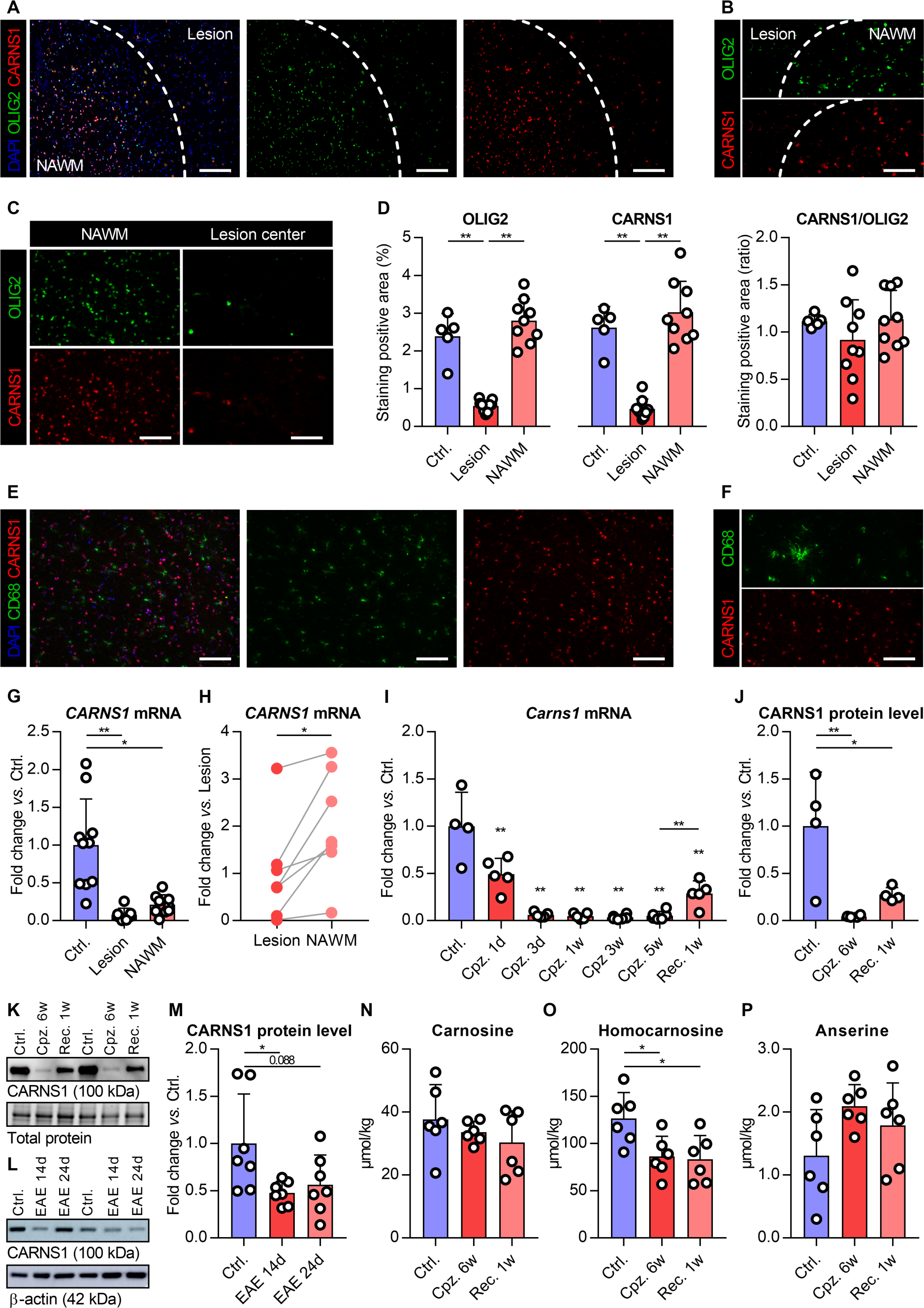
Loss of CARNS1 expression in MS lesions and preclinical MS models. **(a)** Representative images of OLIG2 and CARNS1 immunostaining of a mixed active/inactive MS lesion with surrounding NAWM, **(b)** zoomed in on a lesion border, **(c)** within the lesion center *vs.* within the NAWM. Panels a, b and c are from three different subjects. **(d)** Quantification of immunohistochemistry on human MS lesions and non-demented control tissues. **(e)** Representative images of CD68 and CARNS1 immunostaining: border of a mixed active/inactive lesion, **(f)** zoomed in within lesion border. **(g)** *CARNS1* gene expression in human white matter from non-demented controls, demyelinated lesions and NAWM, including **(h)** paired data from within lesion *vs.* surrounding NAWM. **(i)** *Carns1* gene expression in mouse corpus callosum during cuprizone (Cpz.) and recovery (Rec.). **(j)** CARNS1 protein levels in mouse corpus callosum during cuprizone (Cpz.) and recovery (Rec.). Representative image of CARNS1 detection by western blot in **(k)** mouse corpus callosum and **(l)** spinal cord. **(m)** CARNS1 protein levels in mouse spinal cord during EAE. UHPLC-MS/MS-based detection of **(n)** carnosine, **(o)** homocarnosine and **(p)** anserine in mouse corpus callosum during cuprizone. Scale bars are 200 µm (a), 100 µm (b, c, e, f). Data are mean ± SD. *p<0.05, **p<0.01. NAWM, normal-appearing white matter.

### Immunohistochemistry on mouse spinal cord (EAE) and brain (cuprizone)

Longitudinal spinal cord sections (10 µm) from EAE and control mice were thawed and dried, fixated in acetone (10 min), washed with PBS or PBS with Triton-X100 (PBS-T), blocked (30 min, protein block X0909, Dako), and stained overnight at 4°C using primary antibodies detecting F4/80, GFAP, CD4, SMI-312, inducible nitric oxide synthase (NOS2) or degraded myelin basic protein (dMBP); see details in **Supplementary Table S2**. Following multiple washes, sections were exposed to complementary secondary antibodies for 1 h at room temperature (Thermo Fisher). Using fluorescent microscopy (Leica Microsystems), the entire spinal cord segment was captured in a series of images. The % positive *vs.* total area of the spinal cord was quantified in ImageJ (NIH) for F4/80, GFAP, SMI-312, NOS2 and dMBP stainings. The number of CD4^+^ T cells was calculated by dividing the entire positive stained area by the average area of one cell, and expressed per mm^2^ tissue.

The same protocol was used to stain F4/80, GFAP, NOS2, and dMBP in coronal sections (10 µm, Bregma −1.5 mm) from cuprizone and control brains that were immediately frozen, and to stain MBP and double-stain OLIG2 with CC1 in brains that were PFA-fixated. Paraffin-embedded brains were deparaffinized (xylene-ethanol series), incubated in heated 10 mM citrate buffer, washed in PBS, blocked in 1:10 diluted protein block (30 min; X0909, Dako), and stained overnight at 4°C using primary antibodies against MBP, IBA1, or OLIG2 and CC1 (**Supplementary Table S2**). The % positive *vs.* total area of the corpus callosum was quantified in ImageJ (NIH) for F4/80, GFAP, NOS2, MBP and dMBP stainings. The number of IBA1^+^ cells was calculated by dividing the entire positive stained area by the average area of one cell, and expressed per mm^2^ tissue. The number of OLIG2^+^ cells, OLIG2^+^CC1^−^ cells and OLIG2^+^CC1^+^ cells in the corpus callosum was manually counted and expressed per mm^2^ tissue, and the % of OLIG2^+^ cells that express CC1 was calculated. DAPI was used to mark nuclei in all stainings. Negative control sections omitting the primary antibodies were included for all immunostainings.

### Western blot on mouse spinal cord (EAE) and corpus callosum (cuprizone)

Tissues were diluted in RIPA buffer (50 mM Tris pH 8.0, 150 mM NaCl, 0.5% sodium deoxycholate, 0.1% SDS, 1% Triton-X100, freshly added protease/phosphatase inhibitors [Roche]), and homogenised using stainless steel beads and a QIAGEN TissueLyser II (shaking 1 min, 30 Hz). Following centrifugation (15 min, 12.000 × g, 4°C), supernatants were stored at −80°C. Pierce™ BCA Protein Assay Kit (Thermo Fisher) was used according to manufacturer’s instructions to determine protein concentrations (read at 570 nm wavelength). To detect CARNS1 protein levels in spinal cord or corpus callosum, 5 µg of protein was diluted in loading buffer solution (63 mM Tris Base pH 6.8, 2% SDS, 10% glycerol, 0.004% Bromophenol Blue, 0.1 M DTT), heated for 4 min at 95°C, and separated in polyacrylamide gels at 100-140 V on ice (4-15%, Mini-PROTEAN TGX, Bio-Rad; or as described previously [12]). To detect protein-bound acrolein in spinal cords, 35 µg protein was used. Next, stain-free gels were imaged following UV exposure to visualise total protein content (ChemiDoc MP Imaging System, Bio-Rad). Alternatively, β-actin was stained after stripping the membrane. Proteins were transferred from the gel to an ethanol-immersed PVDF membrane in transfer buffer (30 min, 25 V, 1.0 A, Trans-Blot Turbo Transfer System, Bio-Rad). Membranes were briefly washed in Tris-buffered saline with 0.1% Tween20 (TBS-T), and blocked for 30 min using 3% milk powder in TBS-T. Following overnight incubation at 4°C with primary antibodies against CARNS1 or acrolein-protein adducts (**Supplementary Table S2**), membranes were washed (3 × 5 min), incubated with secondary HRP-conjugated antibodies for 60-90 min at room temperature, washed again (3 × 5 min), and chemiluminescent images were developed in a ChemiDoc MP Imaging System (Bio-Rad) using Clarity Western ECL substrate (Bio-Rad). Protein bands were quantified with Image Lab 6.1 software (Bio-Rad), and normalized to total protein content.

### Gene expression analysis on mouse spinal cord (EAE)

RNA isolation, cDNA synthesis, qPCR, selection of reference genes, and data analysis were performed as described above for human tissue (primer details in **Supplementary Table S4**).

### UHPLC-MS/MS on mouse spinal cord (EAE) and corpus callosum (cuprizone)

Frozen mouse spinal cords (EAE) were homogenized in extraction solution (ultrapure water with 10 mM HCl containing 5 µM carnosine-d4 as internal standard, 1:19 mg:µL ratio) using the QIAGEN TissueLyser II (1 min, 30 Hz). Following centrifugation (20 min, 3000 × g, 4°C), supernatants were immediately diluted in a 3:1 ratio with ice-cold acetonitrile (−20°C), vortexed, and kept on ice for 15min. After a second centrifugation step (20 min, 3000 × g, 4°C), samples were stored at −80°C until the day of UHPLC-MS/MS analysis. Samples were diluted in 4 volumes of 75:25 acetonitrile:water before injection of 2.5 µL into the Xevo TQ-S MS/MS® system. Standard calibration curves containing all HCDs were prepared in homogenates of spinal cords from *Carns1*-KO mice to account for possible matrix effects. Corpus callosum samples (cuprizone) were homogenised as described above, and underwent either exactly the same UHPLC-MS/MS procedures (see results in **Supplementary Fig S2**), or were analysed as described previously [12] (see results in **Fig 1n, 1o, 1p**).

### Flow cytometry on mouse spleen, lymph nodes, and CNS (EAE)

Immune cells were isolated from the spleen, lymph nodes, and CNS (pooled brain and spinal cord). Single cell suspensions from spleen and lymph nodes were obtained by mechanical transfer through a 70 μm cell strainer (Greiner Bio-One). For CNS tissue, the mechanical dissociation was preceded by enzymatic digestion using collagenase D (Roche Diagnostics GmbH) and DNase I (Roche Diagnostics GmbH), and followed by applying a Percoll gradient (GE Healthcare). After centrifugation, CNS cells from mice sacrificed in the chronic EAE stage were resuspended in FACS buffer and stained with Zombi NIR (15 min), followed by blocking (10% rat serum in PBS, 15 min, 4°C) and an antibody panel detecting CD3, CD4, CD8, CD11b, CD19, CD25, CD45, CD183, CD196, and Ly6C (30 min, **Supplementary Table S2**). Cells derived from the CNS, spleen and lymph nodes from acute EAE mice were resuspended in RPMI-1640 medium (Gibco) with 0.5% Pen-Strep and 10% fetal calf serum (FCS), and plated in a 96-well plate (max. 1×10^6^ cells/well). Following overnight incubation at 37°C and 5% CO_2_, cells were exposed to PMA (20 ng/mL), ionomycin (1 µg/mL), and Golgiplug (2 µg/mL) for 4 h. Next, cells were washed, stained with Zombi NIR, blocked, and stained for extracellular markers CD3, CD4, CD8, CD11b, CD19, CD45, and Ly6C (30 min, **Supplementary Table S2**). Finally, cells were resuspended in Cytofix/Cytoperm^TM^ (BD Biosciences) for 30 min, washed with 1X Perm/Wash^TM^ buffer (BD Biosciences), and exposed to antibodies detecting intracellular markers IL-17, IFN-γ, and FoxP3 (15 min, **Supplementary Table S2**). Flow cytometry was performed on a BD LSRFortessa. Our gating strategy excluded doublets and dead cells. FlowJo 10.7.1 software was used for the analyses. Absolute cell numbers were calculated from the relative expression of cell subsets (%) and the total number of cells counted following cell isolation.

### Transmission electron microscopy on mouse corpus callosum (cuprizone)

Glutaraldehyde-fixated mouse brain samples were post-fixated with 2% osmiumtetroxide in 0.05 M sodium cacodylate buffer (1 h, 4°C). Samples were dehydrated by ascending concentrations of acetone. Next, they were impregnated overnight in a 1:1 mixture of acetone and araldite epoxy resin. The samples were embedded in araldite epoxy resin at 60°C and were cut in slices of 70 nm, perpendicular to the corpus callosum, with a Leica EM UC6 microtome. The slices were transferred to 0.7% formvar-coated copper grids (Aurion). Thereafter, samples were contrasted with 0.5% uranyl acetate and lead citrate using a Leica EM AC20. Analysis was performed with a JEM-1400 Flash transmission electron microscope (Jeol) using an EMSIS Xarosa camera. The g-ratio (ratio of the inner axonal diameter to the total outer diameter) was analysed using ImageJ (NIH).

### *In vitro* oligodendrocyte precursor cell differentiation experiments

Primary mouse oligodendrocyte precursor cells (OPCs) were harvested from mixed glial cultures of newborn (P0-2) C57BL/6J mouse cortices. Experiments were approved by the Ethical Committee on Animal Experiments at Hasselt University. Culture conditions were as described previously [28] and outlined in **Supplementary Methods S1a**. To boost OPC formation, bovine insulin (5 µg/mL, Merck) was added after 7 days in culture. At day 14, OPCs were purified via shake-off and seeded on glass coverslips (**Supplementary Methods S1b**). OPCs were exposed to differentiation medium (**Supplementary Methods S1c**) with or without 10 mM L-Carnosine (Flamma) for 6 days, after which cells were washed repeatedly with PBS and fixated with 4% PFA (30 min). Immunocytochemical staining for MBP and O4 was performed to evaluate OPC differentiation. Briefly, each well was incubated with 1% BSA in 0.1% PBS-T (30 min) for blocking, followed by primary antibodies against MBP and O4 (**Supplementary Table S2**) for 4 h at room temperature. Cells were washed with PBS, incubated with complementary secondary antibodies (Thermo Fisher) for 60 min, stained with DAPI nuclear stain, washed, and coverslips were mounted on a glass slide. Fluorescent images (Leica Microsystems) were captured to assess the staining intensity (IntDen) of MBP relative to O4 (ImageJ, NIH).

### Data analysis

Statistical analyses were conducted in GraphPad Prism v7.04. Two independent groups were compared by t-tests or Mann-Whitney U tests. Multiple groups were compared by one-way ANOVA followed by Dunnett’s or Tukey’s multiple comparison test, or by Kruskal-Wallis and Dunn’s test. **Fig 1h** was analysed by a paired t-test. For **Fig 1i**, multiple comparisons were only made to the control group, and a separate t-test was performed to compare the data from Cpz. 5 w *vs.* Rec. 1w. In the EAE analyses, multiple comparisons were only made to the untreated *Carns1*-KO group (any exceptions are clarified in figure captions). For cuprizone experiments, comparisons between *Carns1*-KO and WT mice, or between carnosine-treated and untreated mice, were compared at each time point separately. Data are presented as mean ± SD unless otherwise specified. *p<0.05, **p<0.01 for two-sided hypothesis testing.

## Results

### Loss of CARNS1 in MS lesions and preclinical MS models

We investigated gene expression and protein levels of CARNS1 in human MS lesions and in 2 preclinical MS models (EAE and cuprizone). In human white matter, CARNS1 was located in OLIG2^+^ cells from the oligodendrocyte lineage, confirming our previous observations [19] and available RNA sequencing databases (e.g. [17, 18]). However, OLIG2 and CARNS1 immunoreactivity were both harshly diminished or even absent in the center of demyelinated (PLP^−^) MS lesions, indicating the simultaneous loss of oligodendrocytes and CARNS1 (**Fig 1a, 1b, 1c, 1d**). The surrounding NAWM contained CARNS1 and OLIG2 levels comparable to white matter from healthy donors (**Fig 1d**). CARNS1 was not observed in GFAP^+^ astrocytes, NF-H^+^ neurons/axons, or CD68^+^ macrophages and microglia (**Fig 1e, 1f**) in or around MS lesions. In line with our observation that CARNS1 resides within oligodendrocytes and is lost upon demyelination, *CARNS1* gene expression was 12.6-fold lower in demyelinated MS lesions that lacked mature oligodendrocytes compared to white matter from matched controls (**Fig 1g**). The NAWM of MS patients exhibited a 4.8-fold lower *CARNS1* expression than white matter from matched controls, and a 5.6-fold higher *CARNS1* expression compared to the lesion center (**Fig 1g, 1h**).

We next studied CARNS1 gene expression and protein levels in mice exposed to cuprizone, a toxin that rapidly induces oligodendrocyte loss and demyelination in the CNS and in particular the caudal corpus callosum. *Carns1* mRNA in the corpus callosum was reduced by half following one day and by 17-fold following three days of cuprizone feeding (**Fig 1i**). This robust reduction remained as long as cuprizone feeding was continued, whilst a 5.5-fold increase in *Carns1* mRNA was noted after one week of spontaneous recovery on a normal diet (**Fig 1i**). *Mbp* mRNA levels followed a similar pattern during de- and remyelination (**Supplementary Fig S1a**). Similar gene expression changes were observed in mice undergoing a long-term cuprizone protocol (**Supplementary Fig S1b, S1c**). At the protein level, 6 weeks of cuprizone feeding induced an almost complete loss of CARNS1, which partially recovered upon 1 week of remyelination (**Fig 1j, 1k**). In the EAE model, which is characterised by CNS inflammation due to the infiltration of myelin-reactive T cells, we previously reported a 2-fold reduction in spinal cord *Carns1* mRNA levels during the acute disease stage [12]. This was now confirmed by western blotting, which showed 2.1-fold lower CARNS1 protein levels in spinal cords from EAE compared to healthy mice (**Fig 1l, 1m**).

Next, we assessed how the observed reductions in CARNS1 affect tissue HCD levels. Lower homocarnosine content was noted following 6 weeks of cuprizone feeding as well as 1 week after withdrawal from cuprizone, whilst carnosine and anserine remained unchanged (**Fig 1n, 1o, 1p**; confirmed in a second independent experiment, see **Supplementary Fig S2**). In EAE mice, we previously found a reduction in spinal cord carnosine (2-fold) and homocarnosine (1.4-fold) concentrations during the (sub)acute disease stages [12].

In summary, MS and preclinical demyelinating MS models induce a loss of oligodendrocytes and CARNS1, which may, along with functional/physiological events (e.g. carbonyl overload), affect HCD levels within the CNS during pathology.

### MS patients have normal HCD levels in plasma and CSF

In addition to local (CNS- or lesion-specific) alterations in CARNS1 and HCD levels, we evaluated a potential systemic disturbance of HCD metabolism in people with MS via a targeted UHPLC-MS/MS approach. No differences in circulating carnosine, homocarnosine, N-acetylcarnosine, or balenine were observed between people with MS and age-matched healthy donors (**Fig 2a, 2b, 2c, 2d**). Plasma anserine levels were significantly reduced in people with MS (**Fig 2e, Supplementary Fig S3**), but this decrease was minimal (Δ=-2.2 nM, p=0.03). Follow-up analyses revealed limited or no impact of sex, MS type, and treatment status at the time of blood sampling on plasma HCDs. Hence, measuring circulating HCDs in MS conveys questionable clinical relevance. Carnosine, homocarnosine and N-acetylcarnosine concentrations in CSF samples from people with MS and people without an MS diagnosis did not differ (**Fig 2f, 2g, 2h**). Anserine and balenine were undetectable in CSF. Finally, we assessed the activity of the CN1 enzyme, which hydrolyses circulating (homo)carnosine into its constituent amino acids [34, 35]. CN1 activity was similar in plasma from healthy people and people with MS (**Fig 2i**).

**Fig. 2.**
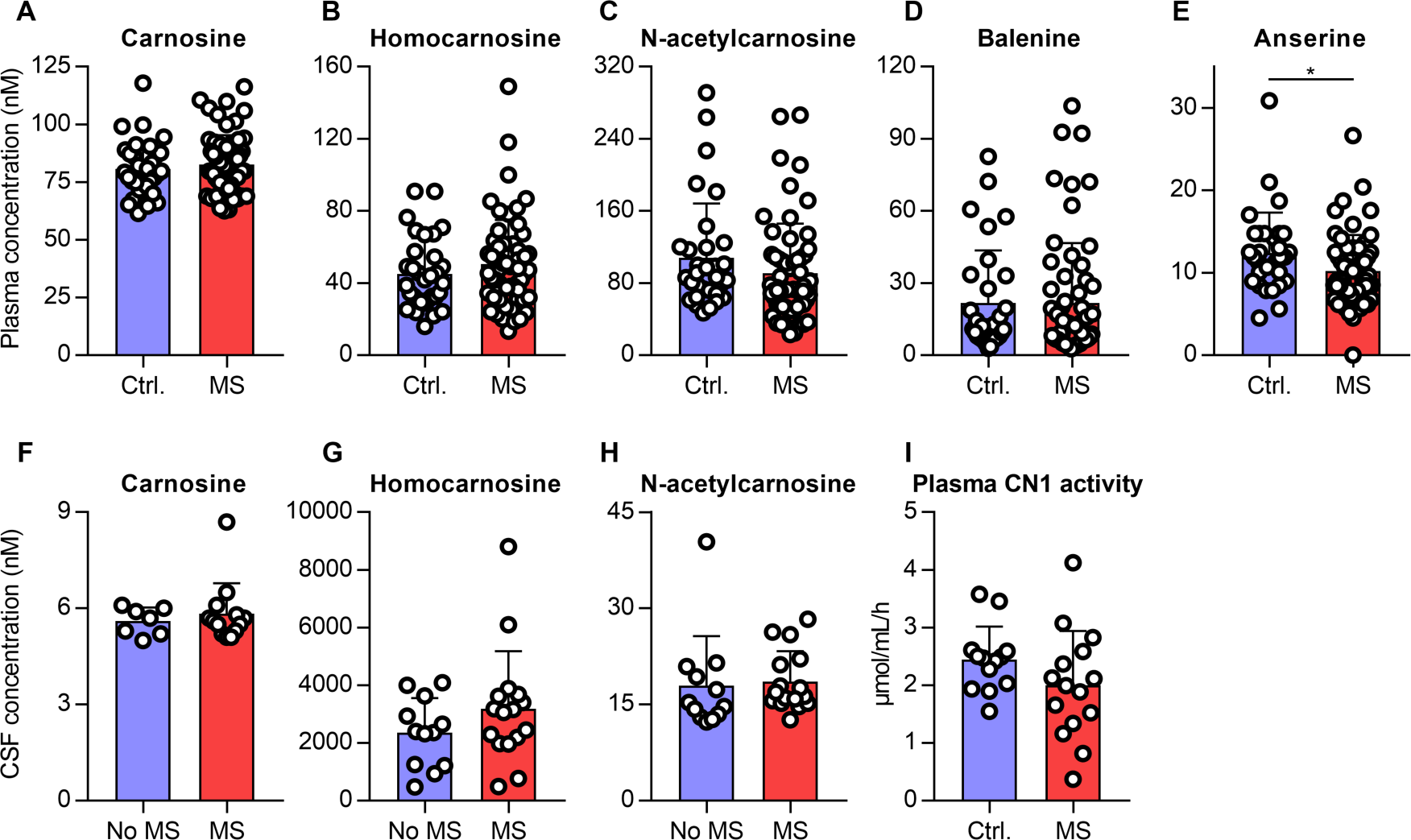
MS patients have normal plasma and CSF histidine-containing dipeptide levels. UHPLC-MS/MS-based detection of **(a-e)** histidine-containing dipeptides in plasma from MS patients and healthy donors, or **(f-h)** CSF from MS patients and people without MS diagnosis. For CSF carnosine, n=8 samples are omitted because carnosine levels were below the LOD of 5 nM. **(i)** Plasma CN1 enzyme activity in MS patients and healthy donors. See also Supplementary Fig S3. Data are mean ± SD. *p<0.05. CSF, cerebrospinal fluid.

### *Carns1* deficiency worsens EAE outcomes and is partially rescued by exogenous carnosine treatment

To mimic the loss of CARNS1 in MS lesions and to study how the loss of CARNS1 affects the immune response, CNS inflammation, and de- and remyelination, we applied EAE and cuprizone models in *Carns1*-KO and WT mice. *Carns1*-KO mice are devoid of endogenous HCDs [19, 32]. In the EAE experiment, WT mice, *Carns1*-KO mice, and *Carns1*-KO mice that received exogenous carnosine treatment (3% in drinking water) were actively immunized and evaluated daily for the degree of tail/hindlimb paresis (**Supplementary Fig S4a**). Compared to WT mice, *Carns1*-KO mice displayed more severe EAE disease scores and an overall higher disease burden (p=0.03, cumulative score 55.6 ± 11.2 *vs.* 27.5 ± 5.4, SEM, **Fig 3a**). The carnosine-treated *Carns1*-KO mice had an intermediate disease burden (cumulative score 41.8 ± 10.4) that was not significantly different from either the WT mice (p=0.41) or the untreated *Carns1*-KO mice (p=0.48). Onset of clinical symptoms was slightly accelerated in *Carns1-* KO mice (p=0.016 *vs.* WT mice, **Fig 3a**). Body weight responses are shown in **Supplementary Fig S4b**. Oral carnosine intake by *Carns1*-KO EAE mice increased spinal cord carnosine content to 3.8-fold higher than WT EAE mice (**Fig 3b**), did not affect homocarnosine levels (**Fig 3c**), and slightly elevated anserine levels (**Fig 3d**).

**Fig. 3.**
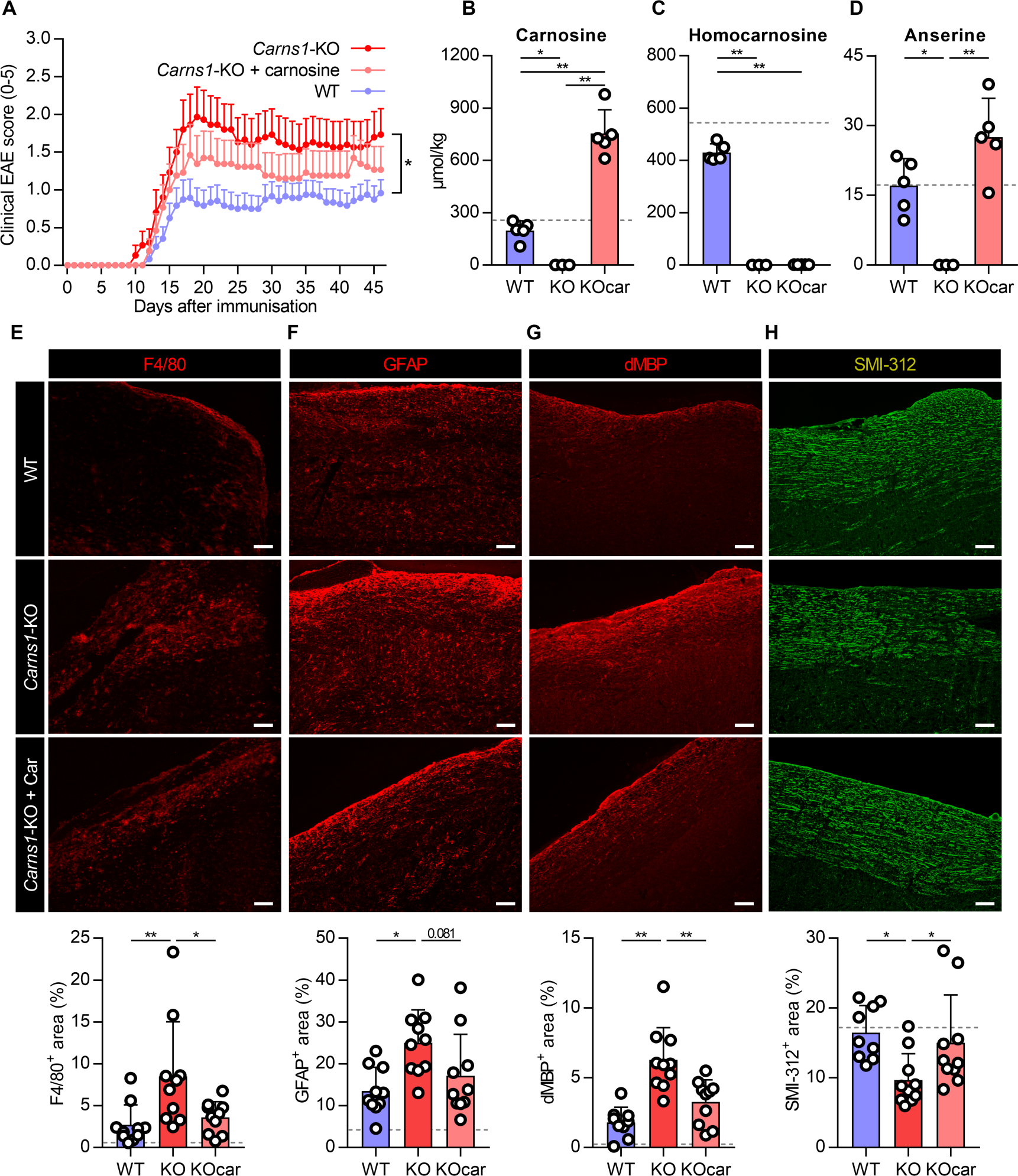
*Carns1* deficiency worsens EAE outcomes and is partially rescued by exogenous carnosine treatment. **(a)** Clinical EAE disease severity, scored daily by a blinded assessor, mean ± SEM, n=13-24/group. UHPLC-MS/MS-based detection of **(b)** carnosine, **(c)** homocarnosine and **(d)** anserine in mouse spinal cord during chronic stage EAE. The dotted line represents healthy control WT mice. Representative images and quantification of mouse spinal cord immunostainings against **(e)** F4/80, **(f)** GFAP, **(g)** degraded MBP, and **(h)** SMI-312 during chronic stage EAE. The dotted line represents the average of healthy control WT and KO mice. Scale bars are 100 µm. Data are mean ± SD. *p<0.05, **p<0.01. For panels b, c and d, multiple comparisons were made between all groups.

By post-mortem histopathological analyses we sought to confirm that *Carns1* deficiency aggravates EAE severity and is partially rescued by exogenous carnosine treatment. An increased lesion load was observed in spinal cords from *Carns1*-KO EAE mice compared to WT EAE mice, as evidenced by greater amounts of F4/80^+^ macrophages/microglia (∼3.1-fold, **Fig 3e**), GFAP^+^ reactive astrocytes (∼1.9-fold, **Fig 3f**), and dMBP^+^ myelin degradation (∼3.5-fold, **Fig 3g**). Neuro-axonal loss (SMI-312 immunoreactivity) was exacerbated in *Carns1*-KO mice (**Fig 3h**). Carnosine treatment was able to fully restore these to WT levels (**Fig 3e, 3f, 3g, 3h**). The number of CD4^+^ T cells in the spinal cord was slightly but not significantly higher in *Carns1*-KO *vs.* WT mice, and was unaffected by carnosine treatment (**Fig 4a**). Using a flow cytometry panel mainly consisting of T cell markers, similar non-significant effects were observed for CD4^+^ T cells, CD4^+^/IFN-γ^+^ T cells, and Ly6C^hi^ inflammatory monocytes (disease peak, **Fig 4b, 4c, 4d, Supplementary Fig S5**) or CD4^+^/CD25^lo^/CXCR3^+^ T cells (chronic stage, **Fig 4e, Supplementary Fig S5**). Phenotyping of immune cells collected from the spleen or lymph nodes at disease peak did not reveal evidence for a peripheral immune-modulatory effect (**Fig 4f, 4g, 4h, 4i, Supplementary Fig S6**). Cytokine production by these cells also did not differ between genotypes (**Supplementary Fig S6**). In contrast, altered inflammatory activity was found within the CNS of *Carns1*-KO mice. Spinal cords from *Carns1*-KO mice had elevated NOS2 protein levels (∼4.2-fold, **Fig 5a**), *Tnf* gene expression (but not *Il1b* or *Il6*, **Fig 5b, 5c, 5d**), and lower *Bdnf* gene expression (**Fig 5e**). This may indicate a skewing of microglia/macrophages towards a more inflammation-promoting phenotype. Genes related to carnosine metabolism are presented in **Supplementary Fig S7**. Carnosine treatment restored NOS2 and *Tnf* expression to WT levels (**Fig 5a, 5b**). This is in line with our previous work showing an anti-inflammatory effect of carnosine treatment in WT mice, and may originate from an impaired ability of *Carns1*-KO mice to quench and eliminate acrolein, a toxic and pro-inflammatory reactive carbonyl that is abundantly generated in EAE and MS lesions [12, 36]. Therefore, we determined acrolein-protein adduct formation by western blotting. This revealed that spinal cords from *Carns1*-KO mice, but not *Carns1*-KO mice that received carnosine treatment, had exaggerated acrolein damage (∼1.9-fold higher *vs.* WT, **Fig 5f**).

**Fig. 4.**
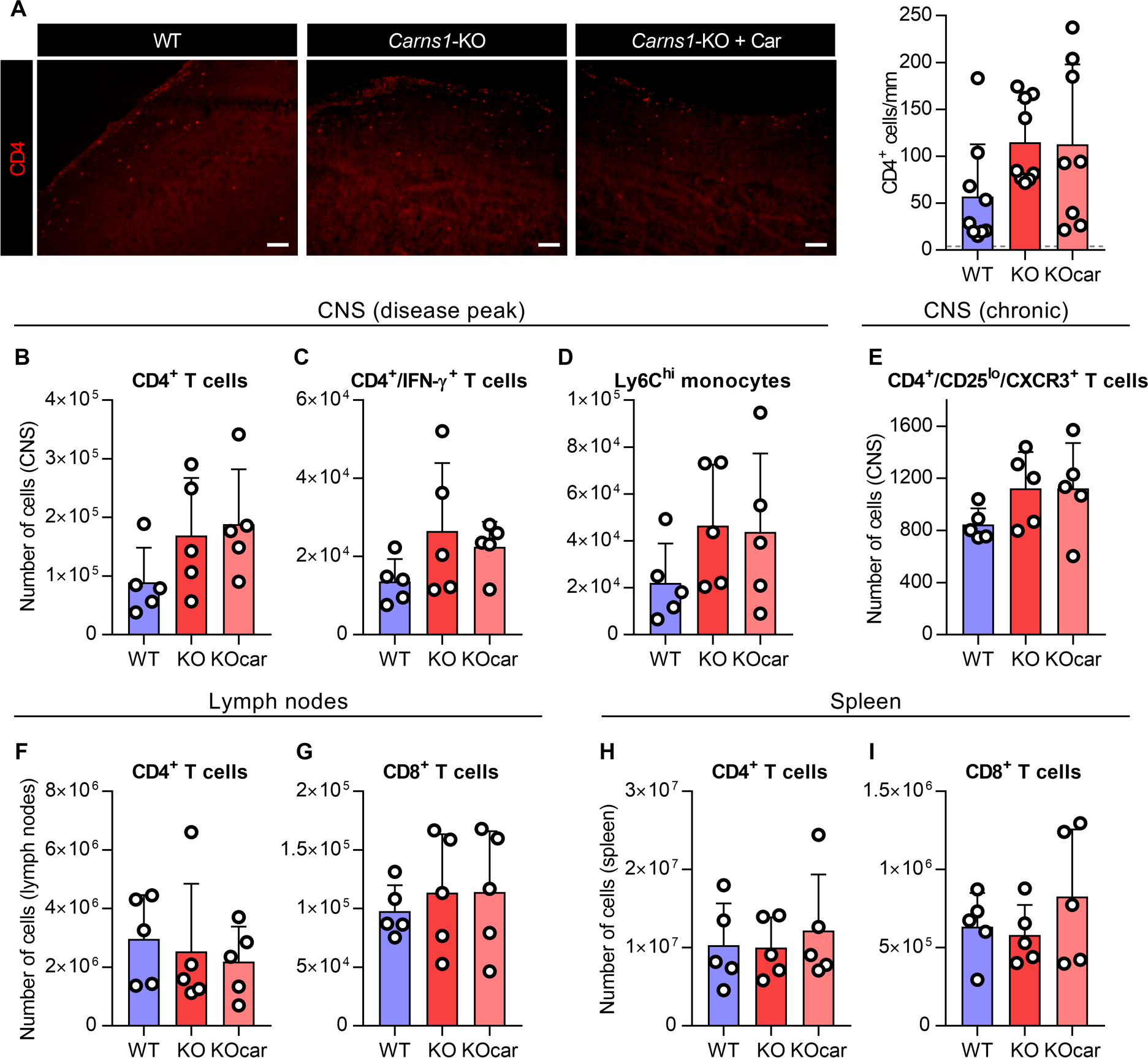
*Carns1* deficiency does not worsen EAE via peripheral immunomodulation. **(a)** Representative images and quantification of mouse spinal cord immunostainings against CD4 during chronic stage EAE. The dotted line represents the average of healthy control WT and KO mice. Flow cytometry analysis of CNS (pooled brain + spinal cord) from EAE mice **(b-d)** at disease peak or **(e)** during the chronic stage. Flow cytometry analysis of **(f, g)** lymph nodes or **(h, i)** spleen from EAE mice at disease peak. Scale bars are 100 µm. Data are mean ± SD. CNS, central nervous system.

**Fig. 5.**
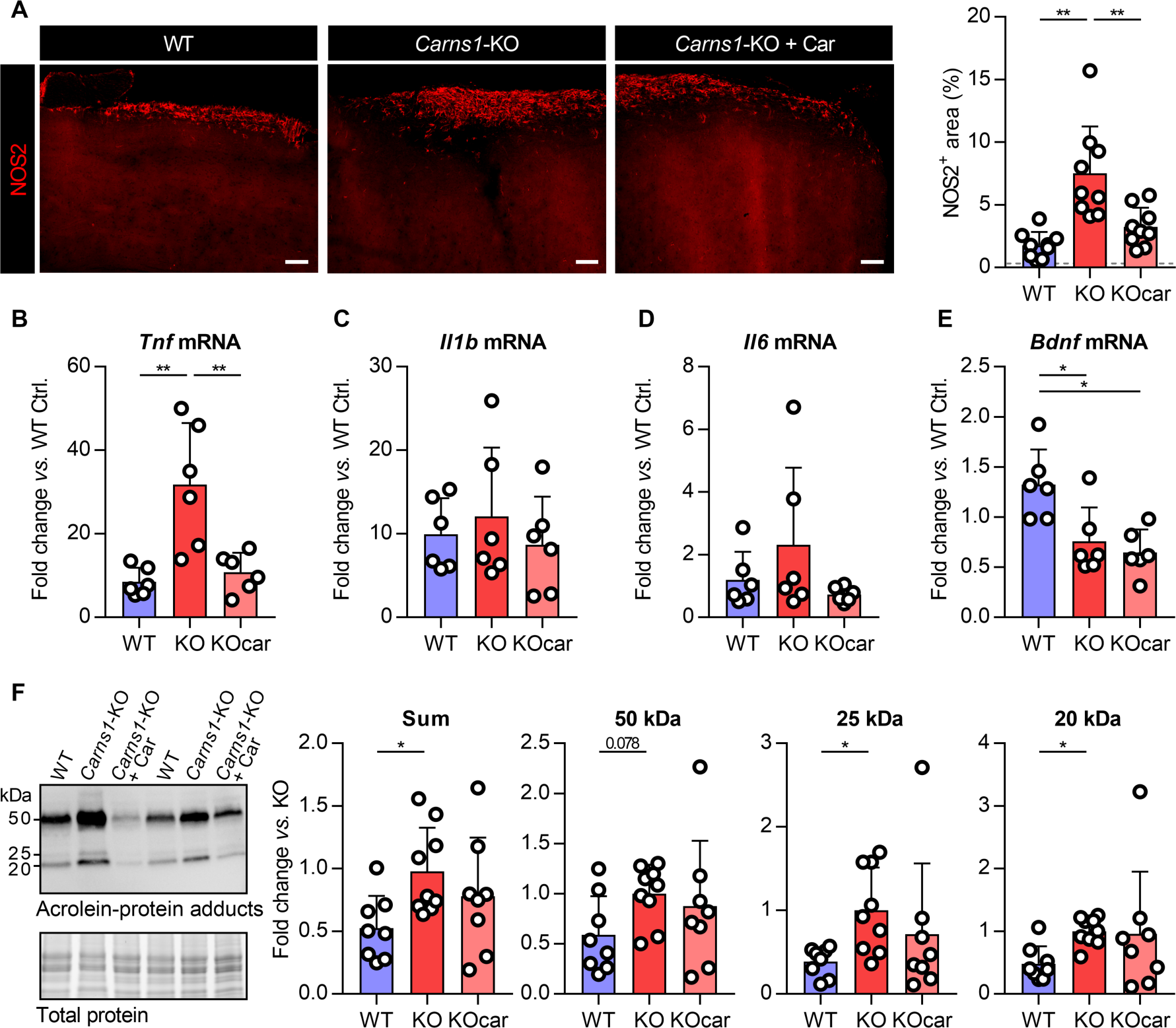
Aggravated neuroinflammatory profile and acrolein damage in *Carns1*-knockout EAE mice. **(a)** Representative images and quantification of mouse spinal cord immunostainings against NOS2 during chronic stage EAE. The dotted line represents the average of healthy control WT and KO mice. **(b-e)** Gene expression analysis of spinal cords during chronic stage EAE. **(f)** Representative western blot image and quantification of acrolein-protein adduct levels in spinal cord during chronic stage EAE. Scale bars are 100 µm. Data are mean ± SD. *p<0.05, **p<0.01. For panel e, multiple comparisons were made between all groups.

Collectively, we show that *Carns1* deficiency aggravates clinical symptoms and pathology within spinal cords from EAE mice. This is likely linked to impaired acrolein quenching and a more pro-inflammatory environment, which could be mitigated via exogenous carnosine treatment.

### *Carns1* deficiency does not affect de- or remyelination in the cuprizone model

Given that CARNS1 in both MS and control subjects is uniquely expressed by cells from the oligodendrocyte lineage, especially as they reach the mature myelinating stage [17, 18], we reasoned that a lack of *Carns1* may accelerate demyelination and hamper remyelination potential of the CNS. To this end, *Carns1*-KO and WT mice were subjected to cuprizone (or control) feeding to induce demyelination, followed by spontaneous recovery (**Supplementary Fig S8a**). Body weight responses were comparable in both genotypes (**Supplementary Fig S8b**). Assessing myelination of the corpus callosum by MBP immunohistochemistry revealed similar degrees of demyelination and remyelination in *Carns1*-KO and WT mice (**Fig 6a**). The same result was obtained for degraded MBP (**Fig 6b**). In addition, *Carns1* deficiency did not affect the number of oligodendrocytes or the oligodendrocyte maturation status following de- or remyelination (**Fig 6c, 6d**). Similarly, g-ratios of corpus callosum axons did not differ between genotypes (**Fig 6e**). Responses of macrophage/microglia (F4/80^+^, **Supplementary Fig S8c**) and astrocyte (GFAP^+^, **Supplementary Fig S8d**) abundance were similar in *Carns1*-KO and WT mice. The NOS2^+^ area was unaltered as well, indicating no effect of genotype on pro-inflammatory activity (**Supplementary Fig S8e**).

**Fig. 6.**
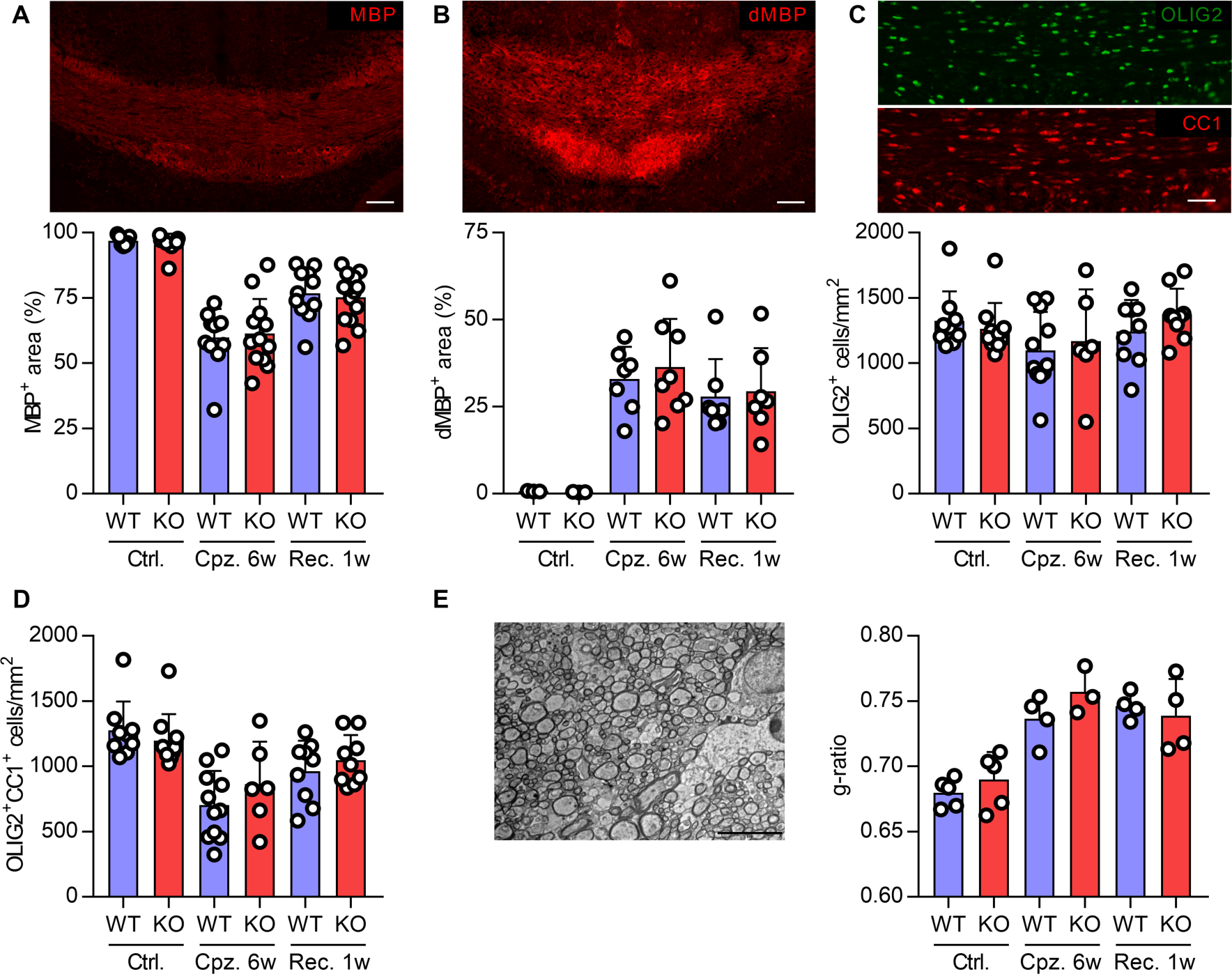
*Carns1* deficiency does not affect de- or remyelination in the cuprizone model. Representative images and quantification of mouse corpus callosum immunostainings against **(a)** MBP, **(b)** degraded MBP, and **(c, d)** OLIG2 and CC1 during cuprizone. **(e)** Representative electron microscopy image and quantification of g-ratio in mouse corpus callosum during cuprizone. Scale bars are 100 µm (a, b), 50 µm (c), 5 µm (e). Data are mean ± SD. Cpz., cuprizone; Rec., recovery.

In summary, CARNS1 expression and the presence of endogenous HCDs do not protect against cuprizone-induced demyelination, and are not required for normal oligodendrocyte development and (re)myelination to occur.

### Exogenous carnosine treatment does not affect de- and remyelination

We also investigated whether exogenous carnosine treatment is a therapeutic agent to limit cuprizone-induced demyelination and/or boost remyelination in WT mice. For this purpose, the same carnosine dosage (3%) that was used in the *Carns1*-KO EAE experiment and that previously improved EAE outcomes in WT mice [12] was administered. Carnosine treatment for 6 weeks led to a 3.3-fold increase in corpus callosum carnosine levels (**Fig 7a**) and a 2-fold increase in anserine levels (**Supplementary Fig S9b**) during cuprizone-induced demyelination. Homocarnosine was unchanged (**Supplementary Fig S9c**). To assess the effect of carnosine loading on remyelination, mice received carnosine during the last 7 days prior to cuprizone withdrawal and during the subsequent 7-day recovery period itself. Using this protocol, the corpus callosum carnosine levels were increased during the entire recovery period to the same extent as in mice treated during the demyelination period (**Supplementary Fig S9d**). Body weight data are shown in **Supplementary Fig S9e**. Assessment of demyelination and remyelination (**Fig 7b**), oligodendrocyte numbers and development status (**Fig 7c, Supplementary Fig S9f**), and abundance of microglia/macrophages (**Supplementary Fig S9g**) in the corpus callosum revealed no effects of carnosine treatment. Finally, a direct effect of exogenous carnosine on the differentiation capacity of primary mouse OPCs was also tested *in vitro*. Exposure to carnosine did not accelerate OPC differentiation, quantified as the MBP/O4 ratio, after 6 days (**Fig 7d**).

**Fig. 7.**
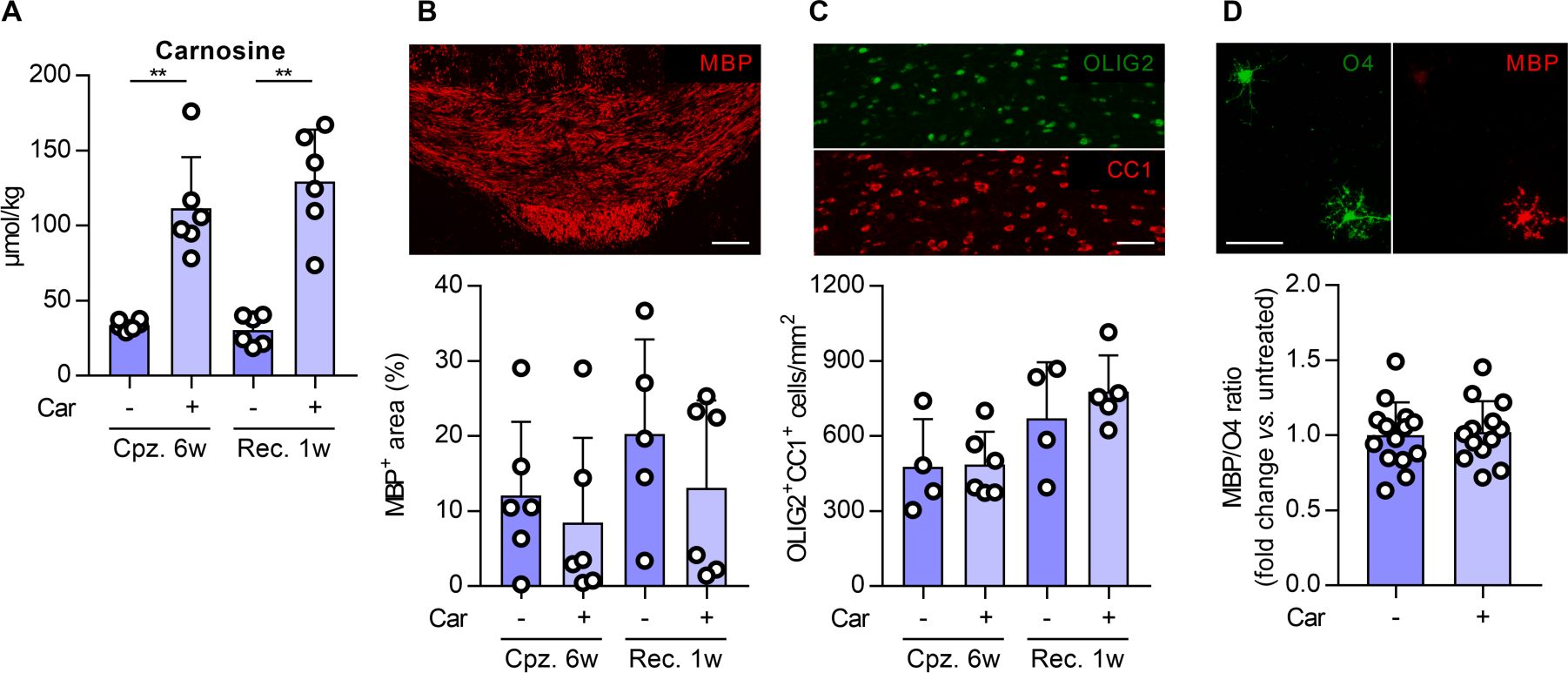
Exogenous carnosine treatment does not affect de- and remyelination. Figures only show treated and untreated cuprizone groups, but exclude healthy control mice and cuprizone mice receiving treatment during the last 7 days of cuprizone feeding (see Supplementary Fig S9a, 9d). **(a)** UHPLC-MS/MS-based detection of carnosine in mouse corpus callosum during cuprizone. Representative images and quantification of mouse corpus callosum immunostainings against **(b)** MBP, and **(c)** OLIG2 and CC1 during cuprizone. **(d)** Representative image and quantification of mouse OPC immunocytochemical stainings against MBP and O4 during differentiation. Scale bars are 100 µm (b, d), 50 µm (c). Data are mean ± SD. **p<0.01. Cpz., cuprizone; OPC, oligodendrocyte precursor cell; Rec., recovery.

In conclusion, carnosine cannot be used as a therapeutic agent to blunt demyelination or accelerate remyelination in the cuprizone model.

## Discussion

CARNS1 is the unique enzyme responsible for (homo)carnosine synthesis in mammals. So far, the role of these dipeptides in the CNS has remained enigmatic, yet their versatile homeostatic properties were hypothesized to provide protection against CNS disease (e.g. via carbonyl quenching). Here we have shown that due to its enrichment within oligodendrocytes of the CNS white matter, CARNS1 is diminished in demyelinated MS lesions. This may have detrimental effects on (the progression of) MS pathology, as the lack of CARNS1 resulted in exacerbated neuroinflammation and acrolein overload in a mouse MS model (EAE). Exogenous carnosine administration partially rescued the disease exacerbation caused by *Carns1* deficiency. Despite its oligodendrocyte-specific localisation, CARNS1 is not required for normal oligodendrocyte development and (re)myelination of the CNS.

In MS, autoreactive lymphocytes invade the CNS parenchyma and trigger neuroinflammation, demyelination, and axonal injury. These CNS-intrinsic events sustain and even progress after immune infiltration wanes [1, 4, 5]. Although the exact pathogenesis of MS is intricate and still under debate [5], compelling evidence supports that oxidative stress and its toxic products (e.g. reactive carbonyls) mediate both acute and chronic CNS disease processes [8, 12, 37–39]. For example, acrolein is abundantly generated in inflammatory lesions from MS patients an EAE mice [12, 37, 40]. To protect against carbonyl overload, the CNS is equipped with an array of mechanisms that neutralize and/or eliminate reactive carbonyls. Carnosine and related HCDs are abundantly present in the CNS (∼500 µmol/kg in mouse spinal cord [19]) and have potent acrolein quenching efficiency [13, 14]. We previously reported that therapeutically increasing spinal cord carnosine levels amplifies acrolein quenching and attenuates CNS inflammation in EAE [12]. In the present report we found that the unique enzyme responsible for carnosine synthesis, CARNS1, is lost upon demyelination in MS. Interestingly, CARNS1 levels dropped more quickly and robustly in cuprizone compared to EAE mice, in contrast to a faster and greater loss of HCDs (especially carnosine) in EAE. This indicates that CARNS1 is not the single determinant of HCD levels and, especially under pathological conditions, functional/physiological events such as binding to acrolein (followed by removal of the conjugate) may contribute to the loss of HCDs during neuroinflammation [12]. Cuprizone mice do not have any carnosine-acrolein conjugate formation in the corpus callosum (unpublished data), which may explain the apparent greater stability of HCDs in this model. In addition, decreasing or increasing carnosine levels in the corpus callosum (via knockout of *Carns1* or carnosine treatment, respectively) do not affect demyelination or remyelination in the cuprizone model. In contrast, our results provide more evidence that carnosine has anti-inflammatory and possibly neuroprotective properties during neuroinflammation, as the deletion of *Carns1* led to profound worsening of EAE disease. The lack of endogenous carnosine in *Carns1*-KO mice resulted in a decreased ability to protect against acrolein-protein adduct formation. It is interesting to note that Wang-Eckhardt *et al.* did not observe an increase in carbonyl-protein adducts in the CNS of healthy aged *Carns1*-KO mice [41], implying that the dependence on HCDs to counter carbonyl overload only manifests under more challenging (pathological) conditions such as EAE. That oligodendrocytes and OPCs are particularly vulnerable to oxidative and carbonyl stress [11] suggests that a vicious cycle may emerge, in which carbonyl overload causes oligodendroglial injury, diminished CARNS1 expression and reduced HCD production; culminating in a weakened protection of the CNS against carbonyl stress, inflammation, demyelination, and neurodegeneration.

Replenishing the spinal cord carnosine pool via exogenous carnosine intake partially rescued the knockout phenotype in the EAE model. Given the unmet need for MS therapeutics that easily cross the blood-brain barrier and exert their effect within the CNS instead of the periphery [4, 42, 43], carnosine or related molecules appear as interesting candidates to temper disease progression in early stages of the disease. Our data indicate that the effects occurred without involvement of peripheral immune modulation, as immune cell profiling in the spleen and lymph nodes revealed no effects from *Carns1* deficiency or carnosine treatment. Within the CNS, however, the absence of HCDs led to larger lesions with exaggerated neuroinflammation, demyelination, and neuro-axonal damage, which were efficiently countered by exogenous carnosine treatment. Previous *in vivo* and *in vitro* experiments from our laboratories [12] and others [44–46] have demonstrated that supplying HCDs (especially carnosine) to macrophages, microglia or astrocytes dampens their pro-inflammatory phenotype upon cellular stress. Nevertheless, future research is warranted to uncover the exact mechanisms and site of action of carnosine therapy in MS, or neuroinflammation in general. Interestingly, a recent case series study (n=3) showed very preliminary but promising effects of an 8-week, 2 g/day carnosine intervention in people MS [47]. Although these results and especially our work support mechanistic evidence for the beneficial impact of carnosine, therapy efficacy in human trials may be optimized by using more stable (CN1-resistant) carnosine analogs with preserved bioactivity and safety. The development of such compounds is another focus of ongoing research [48–50].

In contrast to the profound effects on neuroinflammation in the EAE model, *Carns1* deficiency did not affect de- or remyelination dynamics in the cuprizone model. Based on the (mature) oligodendrocyte-specific expression of CARNS1 and the substantial increase of CARNS1 during recovery following a cuprizone diet, we hypothesized that this enzyme plays a specific and possibly essential role in OPC differentiation and/or myelination. For example, Beyer *et al.* previously identified several small-molecule metabolites that are drastically upregulated during oligodendrocyte development, some of which – most notably taurine – could enhance OPC differentiation and myelination when added exogenously [51]. However, our *in vitro* and *in vivo* data indicate that carnosine and other HCDs are dispensable for oligodendrocyte development and recovery from a demyelinating insult. This is in line with the fact that healthy *Carns1*-KO mice do not display any phenotype, i.e. they exhibit normal postnatal (CNS) development [32, 41]. Augmenting carnosine levels is not an effective strategy to boost OPC differentiation and remyelination. Collectively, these results also raise the question of what proportion of the HCDs produced by CARNS1 is actually needed or utilized within oligodendrocytes themselves, and what proportion is swiftly transported out of the cell towards its surroundings. Our data convincingly show that CARNS1 expression is restricted to oligodendrocytes, but it appears that the site of action of HCDs likely involves other cells or the intercellular space; hence implying multicellular, paracrine dynamics of HCDs within the CNS.

Our observations confirm previous evidence from untargeted approaches suggesting reduced *CARNS1* gene expression in CNS pathologies that feature demyelination. For instance, *CARNS1* was ranked as the 26^th^ and 9^th^ most downregulated gene in corpus callosum and optic chiasm samples, respectively, from MS patients [26]. Additionally, in peri-lesional NAWM from MS patients, *CARNS1* was identified as the 8^th^ most downregulated gene compared to healthy control white matter [25]. We indeed found reduced *CARNS1* gene expression in the NAWM that surrounds MS lesions, further supporting the notion that cellular (sometimes subclinical) alterations can present beyond the lesion site itself [52–54]. In another study, *CARNS1* was the 4^th^ ranked downregulated gene in the anterior cingulate cortex from people with Lewis body dementia, who displayed a general reduction in myelination-related genes in this area [24]. *CARNS1* also appeared as one of the top differentially expressed genes in a single-nucleus RNA sequencing dataset comparing different types of white matter MS lesions and healthy tissues post-mortem [23]. However, this analysis did not confirm any differences between *CARNS1* mRNA in control tissues and NAWM from MS patients, while reporting high *CARNS1* mRNA in active and chronic inactive lesions, but low *CARNS1* mRNA in remyelinated lesions. This again confirms that *CARNS1* expression is sensitive to local changes in the CNS during MS, but it remains unclear why these results differ from our observations of a strong reduction in *CARNS1* gene (and protein) levels.

The local changes in CARNS1 expression and HCD homeostasis were not reflected by systemic alterations (in plasma or CSF). Therefore, HCDs are probably not useful as biomarkers. The fact that 99.1% of the whole-body HCD pool is located in skeletal muscle tissue, which actively releases HCDs into the circulation [19], may prevent the detection of any CNS-linked changes in plasma samples. Moreover, the highly active CN1 enzyme degrades (homo)carnosine in human plasma [34, 35]. N-acetylcarnosine is the most stable (CN1-resistant) circulating HCD but is barely present in the CNS, rendering its plasma levels minimally indicative of CNS metabolism. In agreement with our results, untargeted plasma metabolomics did not find significant differences in N-acetylcarnosine between control and MS samples [55]. So far, metabolomics have not provided evidence for altered HCDs in plasma or CSF in relation to MS [56]. However, a lower circulating carnosine was recently reported in a group of pediatric MS patients [57].

Although CARNS1 produces both carnosine and homocarnosine, we only assessed whether exogenous treatment with carnosine can rescue MS-specific impairments in *Carns1*-KO mice. The metabolism and physiological properties of homocarnosine and carnosine may differ substantially. For instance, carnosine is a more efficient quencher of several reactive carbonyls, whereas homocarnosine may act as a storage pool for the inhibitory neurotransmitter GABA. Our data indicate that *Carns1* deficiency causes exacerbated neuroinflammation via CNS-intrinsic mechanisms, while excluding peripheral immunomodulation. To completely rule out this possibility, an adoptive transfer EAE experiment using activated *Carns1*-KO T cells in naïve recipient mice, and vice versa, should be performed. Since we are the first to study HCD metabolism in MS, additional research is warranted to fully understand the metabolic routes and physiological roles of different HCDs within the CNS during health and disease.

In conclusion, we demonstrate that HCDs are endogenous protectors against carbonyl overload and neuroinflammation, yet their unique synthesizing enzyme CARNS1 is lost upon demyelination in MS. The exacerbation of EAE disease caused by the lack of *Carns1* could be partially rescued via exogenous carnosine treatment, further supporting the use of carnosine-based therapies for people with neuroinflammatory disorders such as MS. In contrast, neither *Carns1* expression nor carnosine treatment directly impacts OPC differentiation or remyelination within the CNS.

## Funding sources

This research was funded by the Special Research Fund (BOF, Hasselt University, Belgium) and Research Foundation - Flanders (FWO Vlaanderen, Belgium).

## Author contributions

Conception and design: JS, BOE, WD. Materials, experiments and data collection: JS, TVdS, SdJ, AvdWB, AT, HB, SPB, ME, EW, TV, NH. Analysis: JS, BOE, WD. Writing original draft: JS. Writing review: WD. Visualization: JS. Revision and approval of final draft: all authors.

## Compliance with Ethical Standards

The authors declare that they have no conflict of interest.

All animal and human experiments complied with local laws and guidelines, and were approved by the appropriate ethical committees; as referred to in the manuscript.

## Acknowledgements

All human brain material has been collected from donors for or from whom a written informed consent for a brain autopsy and the use of the material and clinical information for research purposes had been obtained by The Netherlands Brain Bank (NBB, Netherlands Institute for Neuroscience, Amsterdam; open access www.brainbank.nl). The experiment protocols and methods used for analysing brain samples were conducted with the approval of the NBB and the Medical Ethical Committee of Hasselt University, and carried out according to institutional guidelines. The technical assistance of Anneke Volkaert (Ghent University) and the support from the University Biobank Limburg (UbiLim, prof. Veerle Somers) are greatly appreciated. We thank Flamma (Flamma Group, Chignolo d’Isola, Italy) for providing L-Carnosine. Supplementary Fig S4a, S8a and S9a were created in BioRender.

## SUPPLEMENTARY MATERIAL

### Supplementary Methods S1. *In vitro* OPC differentiation experiments

#### 1a) Mixed glial cell isolation and culture conditions

Following enzymatic dissociation of mouse pup brains using papain (3 U/mL, diluted in DMEM, 1 mM L-cystein; Sigma-Aldrich) for 20 min, mixed glial cell cultures were maintained in DMEM supplemented with penicillin and streptomycin (50 U/mL and 50 mg/mL, Invitrogen) and 10% heat-inactivated fetal calf serum (FCS; Hyclone). Flasks were coated with poly-l-lysine (PLL, 5 µg/mL, Sigma). Cultures were kept at 37 °C in a humidified atmosphere of 8.5% CO_2_.

#### 1b) Shake-off procedure

On day 14, flasks containing mixed glial cultures were placed on an orbital shaker for 45 min (75 rpm, 37 °C) to detach the microglial layer. After removal of the loose microglia, another shake-off was performed (16 h, 250 rpm), resulting in the detachment of OPCs. The OPC-enriched supernatant was collected, incubated for 20 min on a Petri dish and centrifuged (300 × g, 5 min). OPCs were seeded onto 24-well plates on PLL-coated glass coverslips (2×10^5^ cells/well).

#### 1c) OPC differentiation medium

Differentiation medium was prepared as follows: DMEM medium (Sigma-Aldrich), 0.5% penicillin and streptomycin (Invitrogen), 2% horse serum (Sigma-Aldrich), 0.1 mM putrescin (Sigma-Aldrich), 0.3 mM transferrin (Sigma-Aldrich), 0.02 mM progesterone (Sigma-Aldrich), 0.5 µM triiodothyronine (Sigma-Aldrich), 0.2 µM sodium selenite (Sigma-Aldrich), 0.8 mM bovine insulin (Sigma-Aldrich), 0.5 mM L-thyroxine (Sigma-Aldrich), 2% B27 supplement (in house production as described by Chen *et al.*, PMID: 18471889).

Fifty percent of the medium was refreshed every second day.

**Supplementary Table S1.** Clinical details of human brain tissue for immunohistochemistry.

| **Autopsy** | **Age (y)** | **Gender** | **Post-mortem delay** | **Lesion type** | **MS type** |
| --- | --- | --- | --- | --- | --- |
| 01/316 | 43 | Male | 8 h, 30 min | Mixed active/inactive | SPMS |
| 01/214 | 66 | Female | 6 h, 20 min | Mixed active/inactive | SPMS |
| 03/142 | 53 | Male | 5 h, 30 min | Active | PPMS |
| 03/142 | 53 | Male | 5 h, 30 min | Mixed active/inactive | PPMS |
| 04/172 | 43 | Female | 10 h, 45 min | Mixed active/inactive | SPMS |
| 09/152 | 50 | Male | 9 h, 30 min | Mixed active/inactive | PMS |
| 11/080 | 56 | Female | 8 h, 25 min | Mixed active/inactive | PMS |
| 97/404 | 54 | Female | 7 h, 00 min | Non-plaque white matter | PMS |
| 98/309 | 60 | Female | 8 h, 50 min | Non-plaque white matter | PPMS |
| 00/265 | 69 | Female | 13 h, 20 min | Non-plaque white matter | PMS |
| 96/052 | 68 | Female | 5 h, 45 min | White matter | NDC |
| 96/163 | 69 | Female | 8 h, 30 min | White matter | NDC |

NDC, non-demented control; PMS, progressive multiple sclerosis, not specified; PPMS, primary progressive multiple sclerosis; SPMS, secondary progressive multiple sclerosis. Lesion type was assessed by immunohistochemistry (see methods) and classified as proposed by Kuhlmann *et al.* (PMID: 27988845).

**Supplementary Table S2.** Antibodies used for different applications.

| Antigen | Supplier (Cat. No.) | Host | Application | Dilution | Diluent |
| --- | --- | --- | --- | --- | --- |
| **Human samples (IHC)** |  |  |  |  |  |
| CARNS1 | Sigma (HPA038569) | Rabbit | IHC | 1:100 | PBS with 1% BSA |
| CD68 | Dako (M0814 KP1 clone) | Mouse | IHC | 1:100 | PBS with 10% PB, or<br>PBS with 1% BSA |
| GFAP | Sigma (G3893) | Mouse | IHC | 1:100 | PBS with 1% BSA |
| HLA-DR | Thermo Fisher (14-9956-82) | Mouse | IHC | 1:100 | PBS with 10% PB |
| NeuN | Merck (MAB377) | Mouse | IHC | 1:200 | PBS-T with 10% PB |
| NF-H | Abcam (ab8135) | Rabbit | IHC | 1:200 | PBS with 1% BSA |
| OLIG2 | R&D Systems (AF2418) | Goat | IHC | 1:50 | PBS with 1% BSA |
| PLP | Bio-Rad (MCA839G) | Mouse | IHC | 1:100 | PBS with 10% PB |
| **Mouse samples (WB, IHC, ICC)** |  |  |  |  |  |
| Acrolein-protein adducts | Novus Biologicals (NBP2-59359) | Mouse | WB | 1:2000 | TBS-T with 3% milk |
| CARNS1 | Sigma (HPA038569) | Rabbit | WB | 1:1000 | TBS-T with 3% milk |
| CC1 | Merck (OP80) | Mouse | IHC | 1:50 | PBS-T with 10% PB |
| CD4 | Bio-Rad (MCA4635) | Rat | IHC | 1:100 | PBS |
| Degraded MBP (dMBP) | Merck (AB5864) | Rabbit | IHC | 1:1000 | PBS |
| F4/80 | Bio-Rad (MCA497G) | Rat | IHC | 1:100 | PBS |
| GFAP | Sigma (G3893) | Mouse | IHC | 1:200 | PBS with 10% PB |
| IBA1 | Abcam (ab5076) | Goat | IHC | 1:100 | PBS with 10% PB |
| MBP | Merck (MAB386) | Rat | IHC | 1:250 | PBS |
|  |  |  | ICC | 1:500 | PBS-T with 1% BSA |
| NOS2 | Abcam (ab15323) | Rabbit | IHC | 1:100 | PBS |
| O4 | R&D systems (MAB1326) | Mouse | ICC | 1:1000 | PBS-T with 1% BSA |
| OLIG2 | R&D Systems (AF2418) | Goat | IHC | 1:50 | PBS-T with 10% PB |
| SMI-312 | BioLegend (837904) | Mouse | IHC | 1:250 | PBS |
| **Mouse samples (FC)** |  |  |  |  |  |
| Viability (Zombie NIR) | BioLegend (423105) | NA | FC | 1:1000 | PBS |
| CD3-FITC | BioLegend (100204) | Rat | FC | 1:1000 | FB |
| CD4-PB | BioLegend (100428) | Rat | FC | 1:200 | FB |
| CD8-BV510 | BioLegend (100751) | Rat | FC | 1:200 | FB |
| CD11b-PerCP/Cy5.5 | BioLegend (101228) | Rat | FC | 1:200 | FB |
| CD19-BV650 | BioLegend (115541) | Rat | FC | 1:200 | FB |
| CD25-APC | BioLegend (102011) | Rat | FC | 1:200 | FB |
| CD45-AF700 | BioLegend (103128) | Rat | FC | 1:200 | FB |
| CD183-PE | BioLegend (126505) | Hamster | FC | 1:200 | FB |
| CD196-PE/Cy7 | BioLegend (129815) | Hamster | FC | 1:50 | FB |
| Ly6C-BV785 | BioLegend (128041) | Rat | FC | 1:200 | FB |
| FoxP3-AF647 | BioLegend (126408) | Rat | FC | 1:200 | P/W buffer |
| IL-17-PE/Dazzle594 | BioLegend (506938) | Rat | FC | 1:200 | P/W buffer |
| IFN-γ-PE/Cy7 | BioLegend (505826) | Rat | FC | 1:200 | P/W buffer |

BSA, bovine serum albumin; FB, FACS buffer containing PBS with 2% fetal bovine serum and 0.1% sodium azide; FC, flow cytometry; ICC, immunocytochemistry; IHC, immunohistochemistry; NA, not applicable; PB, protein block (X0909, Dako); PBS, phosphate-buffered saline; PBS-T, PBS with 0.05-0.1% Triton-X100; P/W buffer, 1X Perm/Wash™ buffer (BD Biosciences); TBS-T, tris-buffered saline with 0.1% Tween20; WB, western blot.

**Supplementary Table S3.** Clinical details of human brain tissue for gene expression analysis.

| **Autopsy** | **Age (y)** | **Gender** | **Post-mortem delay** | **MS type** |
| --- | --- | --- | --- | --- |
| 13/092 | 62 | Female | 8 h, 10 min | SPMS |
| 06/160 | 69 | Female | 7 h, 30 min | PPMS |
| 11/120 | 73 | Male | 8 h, 45 min | SPMS |
| 12/113 | 59 | Male | 10 h, 45 min | PPMS |
| 15/022 | 82 | Male | 8 h, 05 min | Unknown |
| 01/329 | 65 | Female | 6 h, 00 min | PPMS |
| 11/110 | 67 | Male | 11 h, 00 min | PRMS |
| 14/006 | 56 | Female | 10 h, 30 min | SPMS |
| 02/192 | 72 | Female | 12 h, 00 min | SPMS |
| 96/052 | 68 | Female | 05 h, 45 min | NDC |
| 97/185 | 61 | Female | 10 h, 15 min | NDC |
| 01/016 | 64 | Female | 8 h, 35 min | NDC |
| 95/019 | 54 | Male | 9 h, 10 min | NDC |
| 96/373 | 70 | Male | 7 h, 30 min | NDC |
| 01/322 | 73 | Female | 13 h, 35 min | NDC |
| 09/051 | 51 | Female | 5 h, 36 min | NDC |
| 99/216 | 69 | Male | 19 h, 16 min | NDC |
| 17/124 | 55 | Female | 7 h, 30 min | NDC |
| 03/254 | 82 | Male | 10 h, 00 min | NDC |

NDC, non-demented control; PPMS, primary progressive multiple sclerosis; PRMS, progressive relapsing multiple sclerosis; SPMS, secondary progressive multiple sclerosis.

**Supplementary Table S4.**
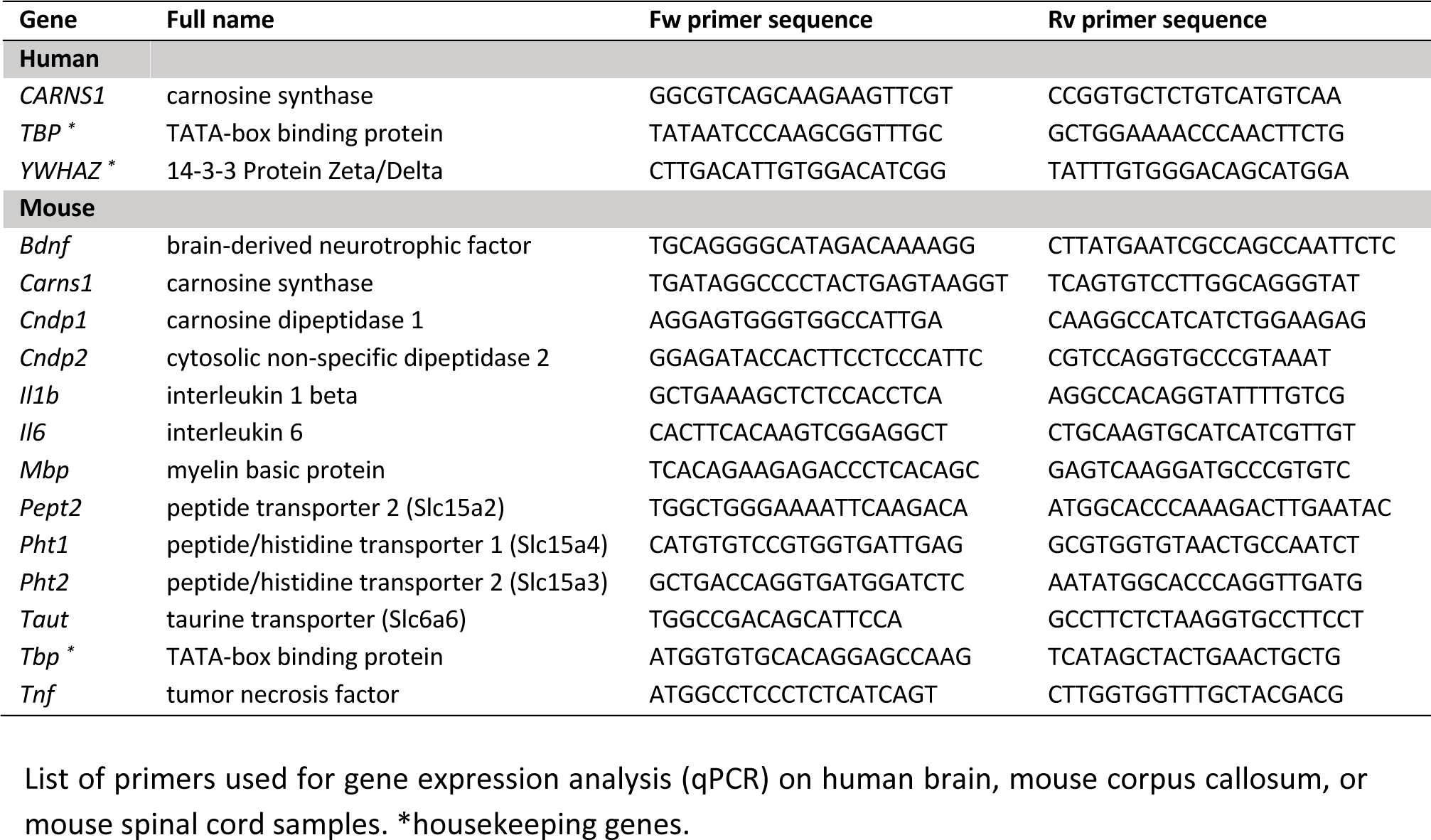
Primer sequences.

**Supplementary Table S5.** Clinical details of blood and CSF donors (UHPLC-MS/MS).

|  | Plasma |  | CSF |  |
| --- | --- | --- | --- | --- |
|  | Control | MS | No MS | MS |
| Number of subjects (n) | 32 | 59 | 12 | 16 |
| Age (y) | 46.4 ± 11.8 | 46.6 ± 11.2 | 48.3 ± 22.7 | 47.3 ± 14.2 |
| Sex (f/m) | 22 / 10 | 45 / 14 | 10 / 2 | 14 / 2 |
| Time since diagnosis (y) | NA | 12.0 ± 10.6 | NA | 6.5 ± 5.3 |
| EDSS | NA | 3.3 ± 2.0 | NA | 4.5 ± 2.8 |
| MS type (RR/SP/PP) | NA | 38 / 19 / 2 | NA | 5 / 7 / 4 |
| Ongoing MS treatment (n) |  |  |  |  |
| Interferon-β | 0 | 13 | 0 | 1 |
| Glatarimer acetate | 0 | 6 | 0 | 0 |
| Alemtuzumab | 0 | 0 | 0 | 1 |
| Natalizumab | 0 | 6 | 0 | 0 |
| Fingolimod | 0 | 8 | 0 | 0 |
| Teriflunomide | 0 | 2 | 0 | 0 |
| Cyclophosphamide | 0 | 9 | 1 | 1 |
| Methotrexate | 0 | 3 | 0 | 0 |
| None | 32 | 12 | 11 | 13 |

EDSS, expanded disability status scale; NA, not applicable; PPMS, primary progressive multiple sclerosis; RRMS, relapsing-remitting multiple sclerosis; SPMS, secondary progressive multiple sclerosis. The 'No MS' group includes people without a diagnosis of MS, but possibly other complaints or diseases.

**Supplementary Table S6.** Clinical details of blood donors (CN1 activity).

|  | **Control** | **MS** |
| --- | --- | --- |
| Number of subjects (n) | 13 | 15 |
| Age (y) | 44.2 ± 14.3 | 49.7 ± 12.5 |
| Sex (f/m) | 10 / 3 | 11 / 4 |
| Time since diagnosis (y) | NA | 5.0 ± 4.6 |
| EDSS | NA | 5.0 ± 2.1 |
| MS type (RR/SP/PP) | NA | 6 / 6 / 3 |
| Ongoing MS treatment (n) |  |  |
| Interferon-β | 0 | 1 |
| Natalizumab | 0 | 1 |
| DMF | 0 | 1 |
| Cyclophosphamide | 0 | 1 |
| None | 13 | 11 |

EDSS, expanded disability status scale; NA, not applicable; PPMS, primary progressive multiple sclerosis; RRMS, relapsing-remitting multiple sclerosis; SPMS, secondary progressive multiple sclerosis.

**Supplementary Fig. S1.**
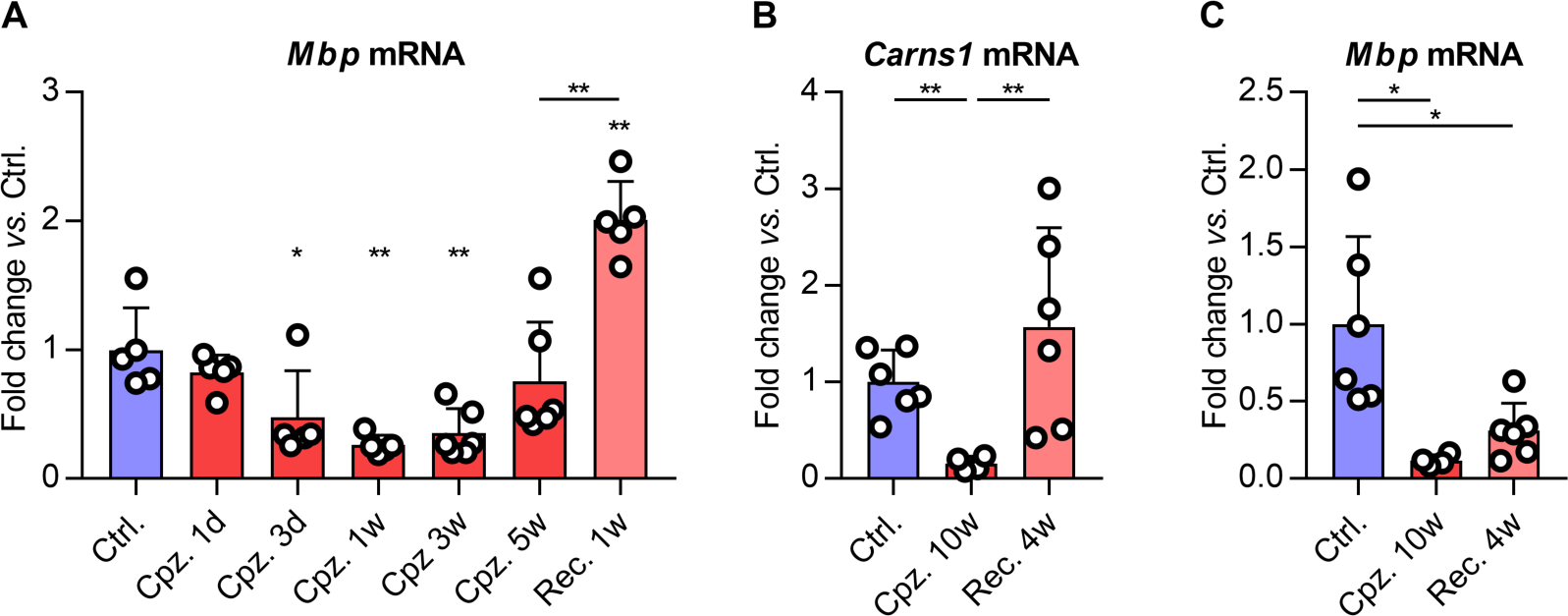
**(a)** *Mbp* gene expression in mouse corpus callosum during cuprizone (Cpz.) and recovery (Rec.). **(b)** *Carns1* and **(c)** *Mbp* gene expression in mouse corpus callosum during a longer-term cuprizone (Cpz.) and recovery (Rec.) experiment. Data are mean ± SD. *p<0.05, **p<0.01.

**Supplementary Fig. S2.**
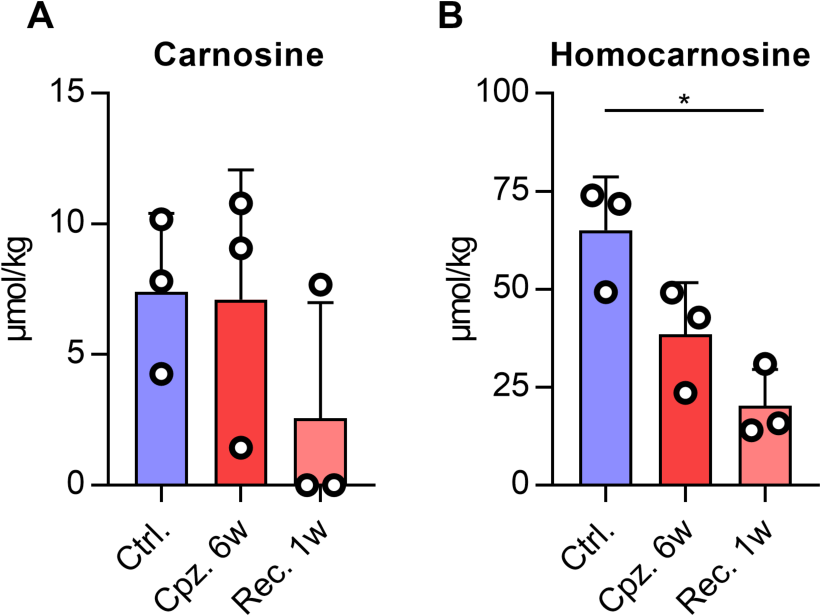
UHPLC-MS/MS-based detection of **(a)** carnosine and **(b)** homocarnosine in mouse corpus callosum during cuprizone. Data are mean ± SD. *p<0.05.

**Supplementary Fig. S3.**
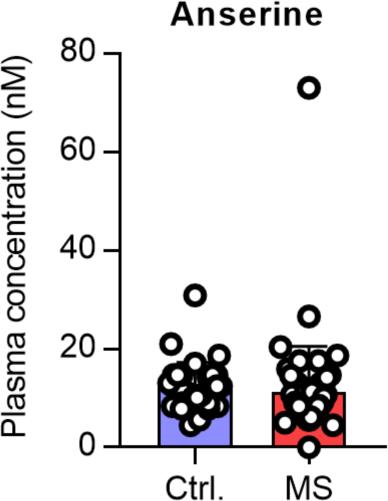
UHPLC-MS/MS-based detection of anserine in plasma from MS patients and healthy donors. One outlier was identified in the MS group and removed from Fig 2e. Data are mean ± SD.

**Supplementary Fig. S4.**
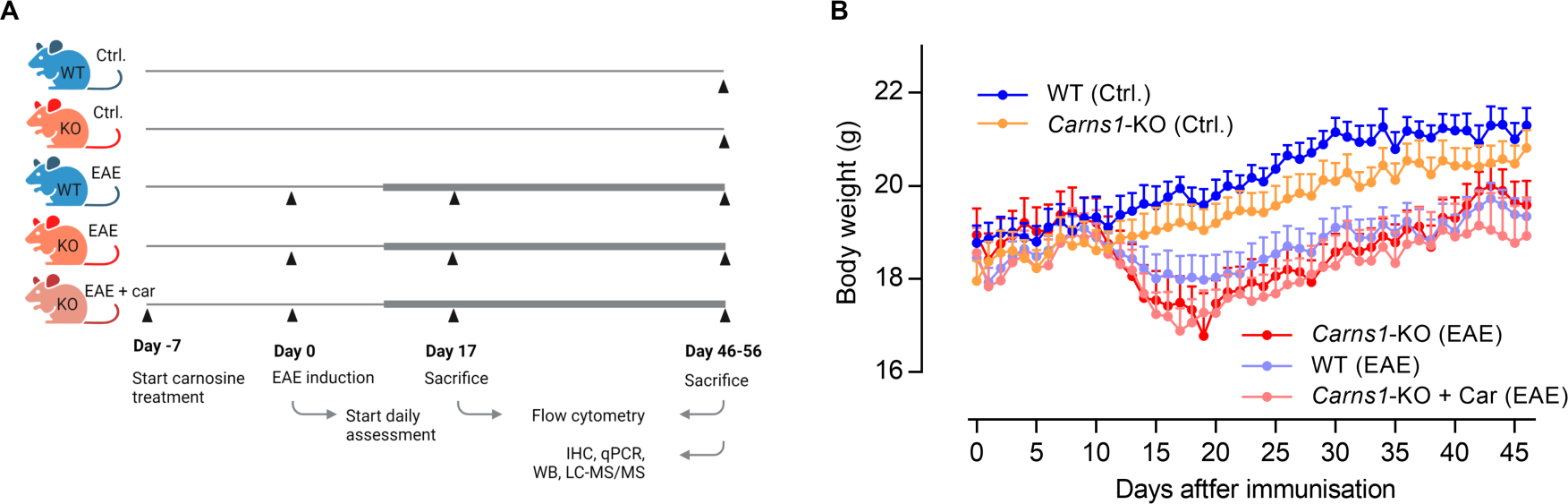
**(a)** Graphical representation of the EAE experiment with *Carns1*-KO and WT mice. A bold line represents clinical disease symptoms in EAE mice. **(b)** Body weight responses. Data are mean ± SEM. Car, 3% carnosine treatment in drinking water. IHC, immunohistochemistry; UHPLC-MS/MS, ultra-high performance liquid chromatography tandem mass spectrometry; qPCR, quantitative polymerase chain reaction; WB, western blot.

**Supplementary Fig. S5.**
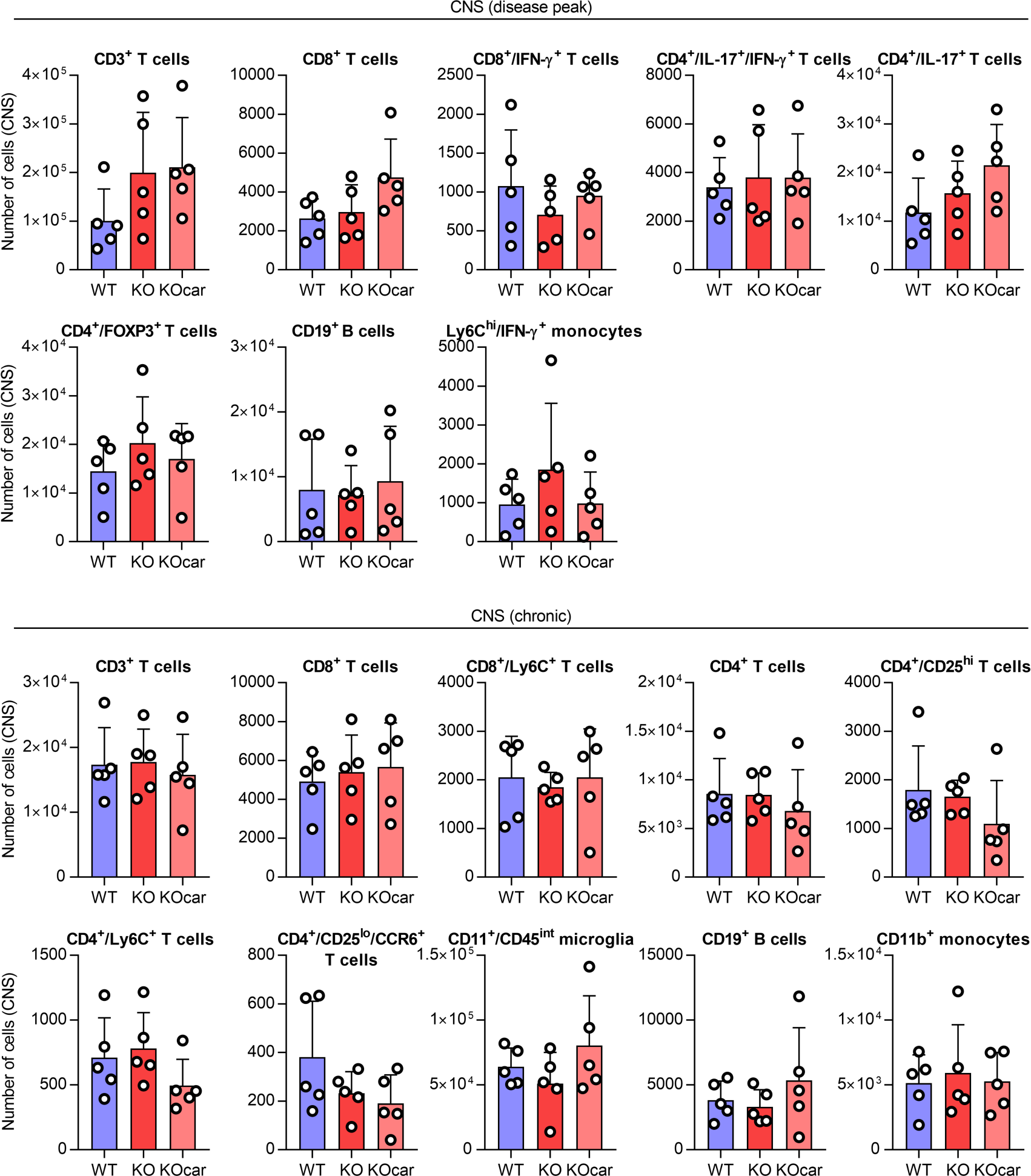
Flow cytometry analysis of CNS (pooled brain + spinal cord) from EAE mice at disease peak or during the chronic stage. Data are mean ± SD.

**Supplementary Fig. S6.**
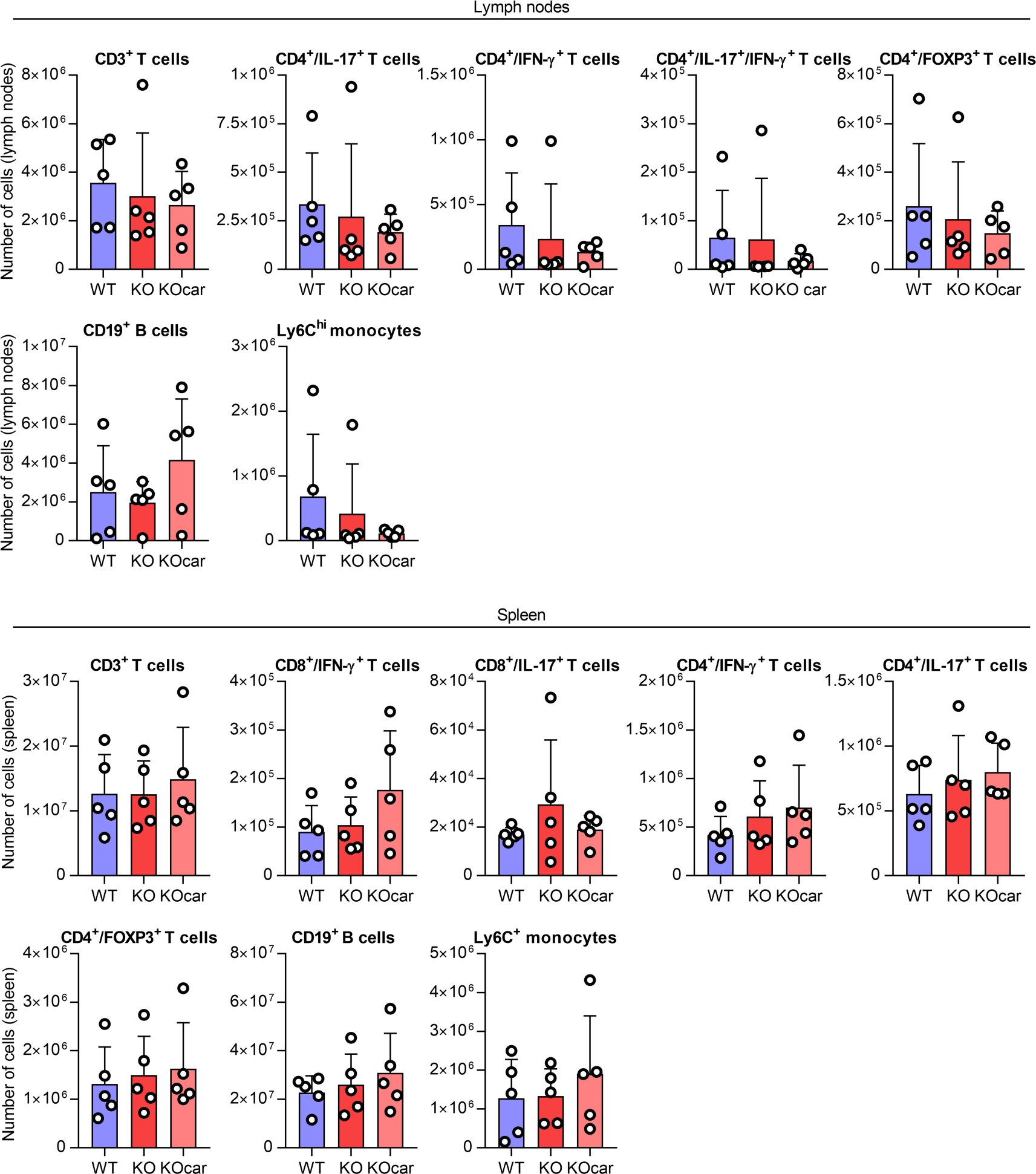
Flow cytometry analysis of lymph nodes or spleen from EAE mice at disease peak. Data are mean ± SD.

**Supplementary Fig. S7.**
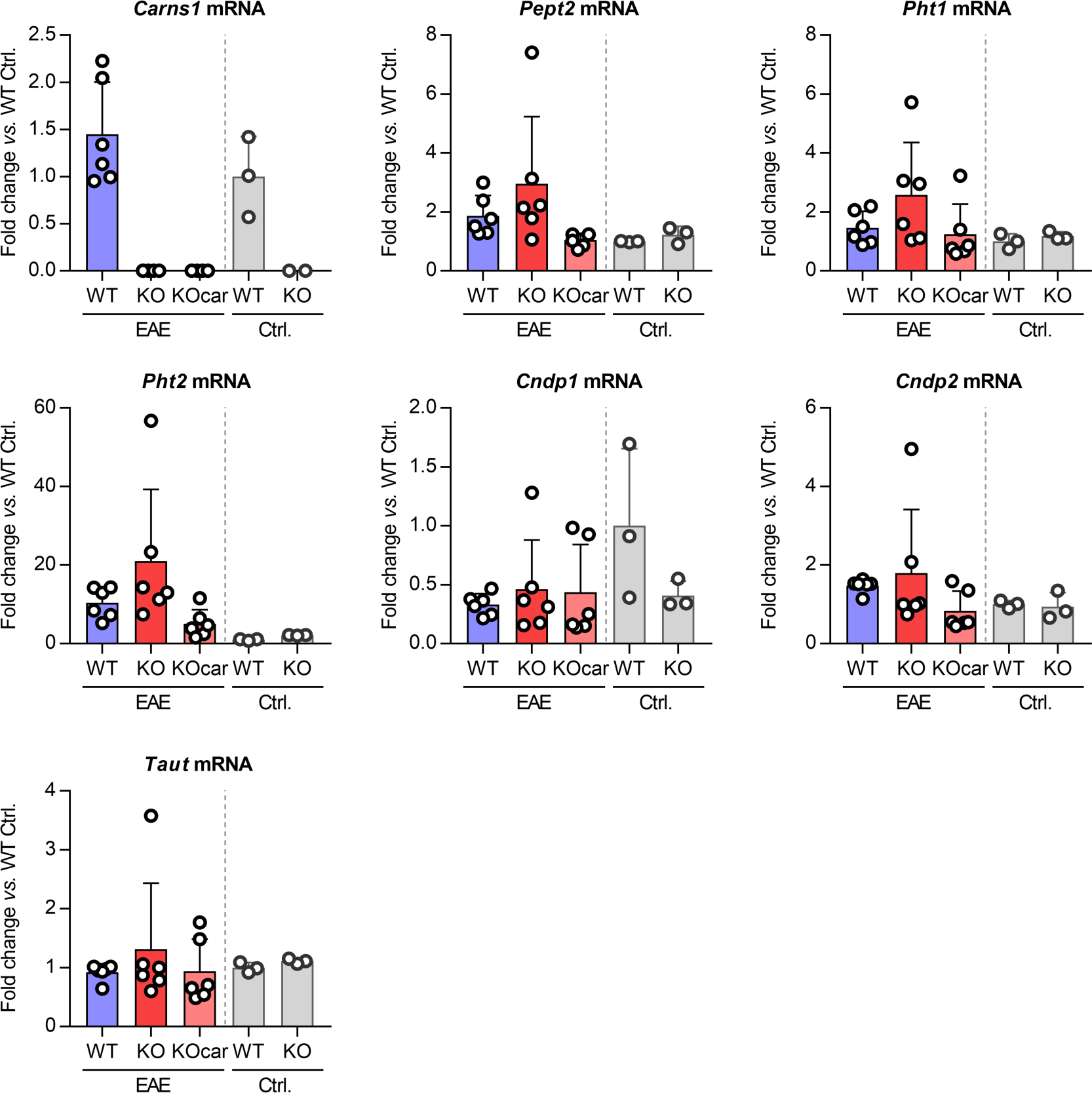
Gene expression analysis of spinal cords during chronic stage EAE. Data are mean ± SD.

**Supplementary Fig. S8.**
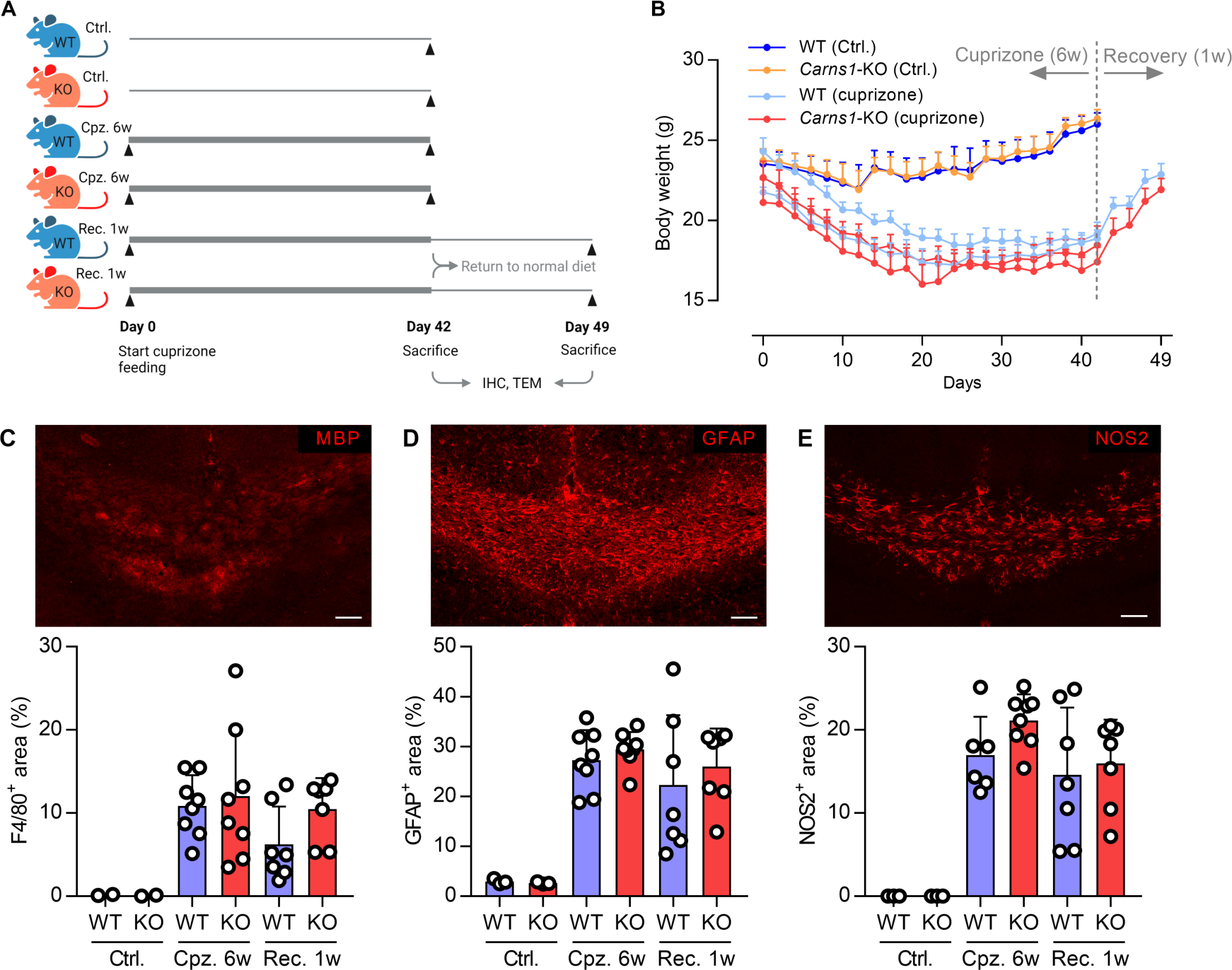
**(a)** Graphical representation of the cuprizone experiment with *Carns1*-KO and WT mice. A bold line represents cuprizone feeding (0.3% w/w). **(b)** Body weight responses. Data are mean ± SEM. Representative images and quantification of mouse corpus callosum immunostainings against **(c)** F4/80, **(d)** NOS2 and **(e)** GFAP during cuprizone. Scale bars are 100 µm. Data are mean ± SD. Cpz., cuprizone; IHC, immunohistochemistry; Rec., recovery; TEM, transmission electron microscopy.

**Supplementary Fig. S9.**
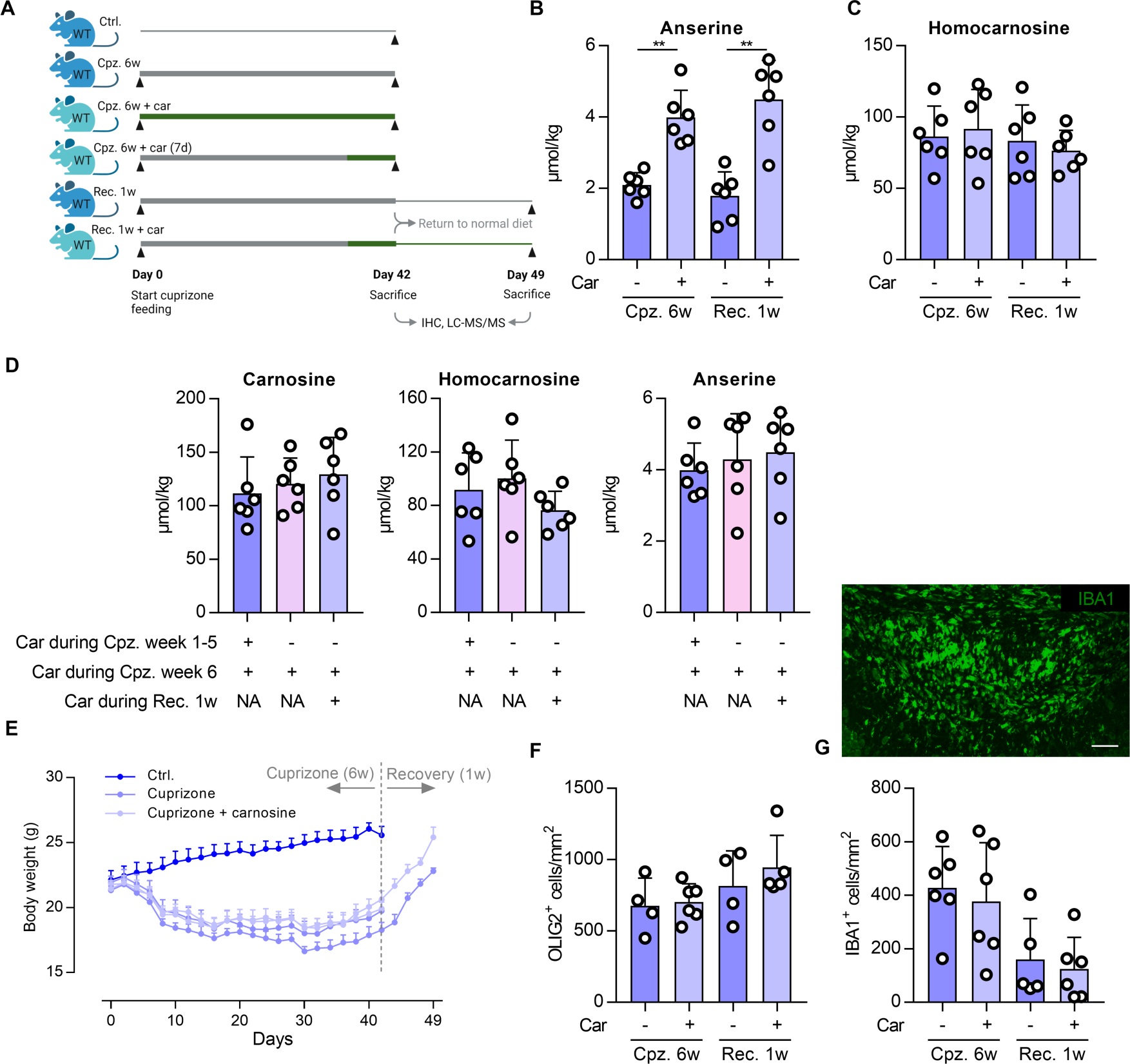
**(a)** Graphical representation of the cuprizone experiment investigating exogenous carnosine treatment in cuprizone mice. A bold line represents cuprizone feeding (0.3% w/w). Green color indicates carnosine treatment. UHPLC-MS/MS-based detection of **(b)** anserine and **(c)** homocarnosine in mouse corpus callosum during cuprizone and recovery with/without carnosine treatment. Data are mean ± SD. **(d)** UHPLC-MS/MS-based detection of carnosine, homocarnosine and anserine in mouse corpus callosum during cuprizone and recovery with/without carnosine treatment. Data are mean ± SD. **(e)** Body weight responses. Data are mean ± SEM. Representative images and/or quantification of mouse corpus callosum immunostainings against **(f)** OLIG2 and **(g)** IBA1 during cuprizone. Scale bar is 100 µm. Data are mean ± SD. Car, 3% carnosine treatment in drinking water; Cpz., cuprizone; IHC, immunohistochemistry; UHPLC-MS/MS, ultra-high performance liquid chromatography tandem mass spectrometry; Rec., recovery.

## Notes

### Competing Interest Statement

The authors have declared no competing interest.

